# A two-pool mechanism of vesicle release in medial habenula terminals underlies GABA_B_ receptor-mediated potentiation

**DOI:** 10.1101/2022.10.28.514202

**Authors:** Peter Koppensteiner, Pradeep Bhandari, Cihan Önal, Carolina Borges-Merjane, Elodie Le Monnier, Yukihiro Nakamura, Tetsushi Sadakata, Makoto Sanbo, Masumi Hirabayashi, Nils Brose, Peter Jonas, Ryuichi Shigemoto

**Author notes:** These authors contributed equally (Peter Koppensteiner, Pradeep Bhandari, Cihan Önal). Corresponding authors: Ryuichi Shigemoto, Peter Koppensteiner.

## Abstract

GABA_B_ receptor (GBR) activation inhibits neurotransmitter release in axon terminals in the brain, except in medial habenula (MHb) terminals, which show robust potentiation. However, mechanisms underlying this enigmatic potentiation remain elusive. Here, we report that GBR activation induces a transition from tonic to phasic release accompanied by a 4-fold increase in readily releasable pool (RRP) size in MHb terminals, mirrored by a similar increase in the docked vesicle number at the presynaptic active zone (AZ). The tonic and phasic release vesicles have distinct coupling distances. We identified two vesicle-associated molecules, synaptoporin and CAPS2, selectively involved in tonic and phasic release, respectively. Synaptoporin mediates augmentation of tonic release and CAPS2 stabilizes readily releasable vesicles during phasic release. A newly developed “Flash and Freeze-fracture” method revealed selective recruitment of CAPS2 to the AZ during phasic release. Thus, we propose a novel two-pool mechanism underlying the GBR-mediated potentiation of release from MHb terminals.

## Introduction

The synaptic connection from the medial habenula (MHb) to the interpeduncular nucleus (IPN) is a phylogenetically conserved pathway involved in emotion- and addiction-related behaviors^1–4^. The most striking peculiarity in this pathway is an enhancement of unparalleled scale of neurotransmitter release from terminals originating from cholinergic neurons in the ventral MHb by activation of presynaptic GABA_B_ receptors (GBRs)^4^, usually inhibitory G-protein coupled receptors^5^. Except for an increase in presynaptic Ca^2+^ influx, it is currently unknown which mechanisms mediate this increase in release. Another unique feature of the MHb-IPN synapse is the exclusive use of Cav2.3 for release^1^. However, Cav2.3-mediated release is not necessarily potentiated by GBR activation since it is inhibited in terminals originating in the dorsal MHb^1,6^.

Across the nervous system, synaptic terminals respond in either one of two fundamental ways to the repeated excitation at high frequency, facilitation of neurotransmitter release called tonic release, or depression of neurotransmitter release called phasic release^7–9^. While the depression of phasic release is mediated by a depletion of the readily releasable pool (RRP) of synaptic vesicles (SVs), the facilitation of tonic release is likely mediated by a progressive increase in release probability (P_r_) by residual Ca^2+^ ^7,10^. Although the molecular properties determining the release modes in which a certain synapse type operates are incompletely understood, relevant factors include Ca^2+^ influx, stimulation frequency and the priming and docking states of SVs^7–9,11^.

Here, we report that, following GBR activation, MHb terminals transition from a facilitating, tonic to a depressing, phasic neurotransmitter release mode at a physiological stimulation frequency. This transition is induced by a GBR-mediated increase in Ca^2+^ influx, which recruits additional SVs to the RRP. Using “Flash and Freeze” in acute IPN slices, we observed an increase in docked SVs following GBR activation to the same extent as the increase in RRP size. We screened SV-associated proteins expressed in MHb neurons and found synaptoporin and CAPS2 selectively involved in tonic and phasic release, augmenting release and retaining readily releasable SVs, respectively. Using a new method for nanoscale visualization of membrane-associated proteins within milliseconds after exocytosis, named “Flash and Freeze-fracture”, we discovered CAPS2 recruitment to the presynaptic active zone (AZ) and its stabilization there during phasic release.

## Results

### Transition from tonic to phasic neurotransmitter release by GBR activation

Rodent MHb neurons are active at ~10 Hz *in vivo^12–14^* but postsynaptic responses in IPN neurons following repetitive stimulation of MHb axons at this physiological frequency have never been studied. Therefore, we recorded rostral/central IPN neurons in whole-cell mode and measured excitatory postsynaptic currents (EPSCs) evoked by electrical stimulation of the fasciculus retroflexus (FR), a fiber bundle arising in the habenula. In order to keep the MHb-IPN pathway intact, we prepared 1 mm-thick angled slices as described previously^1^ (Fig. 1A). Under baseline conditions, 10–Hz stimulation for three seconds produced EPSC responses that increased in amplitude with consecutive stimuli (Fig. 1B). Application of the GBR agonist R(+)-baclofen (1 μM) greatly potentiated initial EPSC amplitudes in the 10–Hz train but continued stimulation progressively reduced subsequent EPSC amplitudes (Fig. 1B-D). Normalization of EPSC amplitudes to the corresponding first baseline EPSC revealed that the enhanced EPSC amplitudes by baclofen decayed to the level of the facilitated baseline EPSC amplitudes (Fig. 1D). This implies that GBR-mediated depressing neurotransmission occurred in addition to and not instead of the baseline facilitating release. Thus, MHb terminals exhibited a facilitating, tonic neurotransmitter release pattern under baseline conditions and transitioned to a depressing, phasic release pattern after GBR activation.

**Figure 1:**
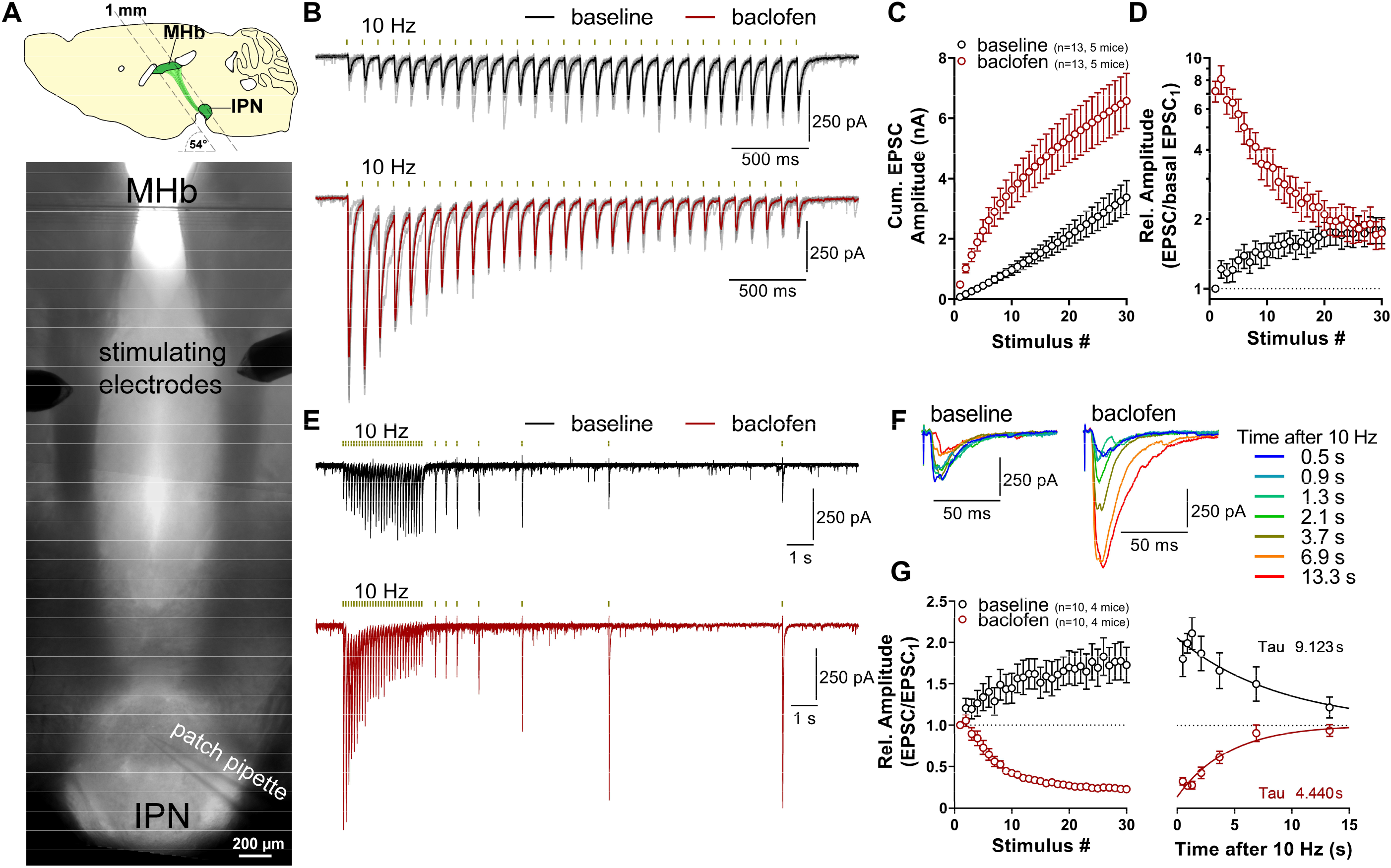
Stimulation of MHb axons at a physiological frequency reveals transition from tonic to phasic neurotransmitter release by baclofen. **A** Scheme of the 1 mm thick angled slice preparation (top) and an example of the recording configuration in the resulting slice (bottom). **B** Example traces of EPSCs evoked by 10–Hz stimulation in one cell before and after the application of baclofen. Greyed responses represent individual sweeps, and bold responses represent the average of the individual traces. **C** Quantification of baseline and baclofen responses using cumulative EPSC amplitudes. **D** Plot of EPSC responses after normalization of EPSC amplitudes to the amplitude of the corresponding first baseline EPSC **E** Overlay of 7 traces of 10 Hz responses at baseline and during baclofen in one cell with single stimulations at varying intervals following the 10 Hz stimulus. **F** Overlay of single stimuli color-coded with stimulation time. **G** Quantification of recovery from tonic release augmentation and phasic release depletion. Using exponential fit, both short-term plasticity exhibited recovery times in the order of seconds. See also Extended Data Fig. 1

Ventral MHb terminals co-release glutamate and acetylcholine^15^, therefore, we tested whether the tonic and phasic release are selectively mediated by one of the two transmitters. Cholinergic EPSC trains in the presence of the AMPA receptor blocker DNQX (10 μM) still transitioned to a phasic response pattern after GBR activation (Extended Data Fig. 1A) similar to that of glutamatergic EPSC trains in the presence of cholinergic blockers (50 μM hexamethonium, 5 μM mecamylamine) (Extended Data Fig. 1B). Although the cholinergic RRP size (0.72 ± 0.18 nA, n=8) was much smaller than the glutamatergic one (3.57 ± 0.47 nA, n=8; Extended Data Fig. 1C), cholinergic P_r_ was not significantly different from that of glutamatergic neurotransmission (Extended Data Fig. 1C). Thus, the GBR-mediated phasic release induction equally affects cholinergic and glutamatergic release, though glutamate mainly contributes to the EPSCs.

We then quantified the recovery from activity-dependent modulations of tonic and phasic release (Fig. 1E, F). The facilitated baseline responses decayed exponentially with a tau of 9.1 s (Fig. 1G), suggesting that this facilitation is augmentation^16^. Similarly, depression of phasic release recovered with a tau of 4.4 s, suggesting that the EPSC decrease results from RRP depletion. In conclusion, the two modes of neurotransmitter release from MHb terminals exhibit distinct short-term plasticity and recover from the activity-dependent modulation within seconds.

### Role of presynaptic Ca^2+^ in the GBR-mediated enhancement of neurotransmitter release

We next explored the role of presynaptic Ca^2+^ in the GBR-mediated transition from tonic to phasic release. Using AAV injections into the MHb of ChAT-IRES-Cre mice, we expressed axon-GCaMP6s in cholinergic neurons. FR stimulation at 10 Hz for three seconds increased presynaptic GCaMP fluorescence in the IPN 4.0 ± 0.6 fold of resting fluorescence intensity whereas application of baclofen further increased it 6.5 ± 0.9 fold (n=5 slices, 5 mice; Fig. 2A, B). However, presynaptic GBRs are known to inhibit the influx of Ca^2+^ through voltage-gated Ca^2+^ channels (VGCCs) and no report shows direct GBR-mediated increases in Ca^2+^ influx for any VGCCs in reconstituted systems^5^. Using simulations based on our Ca^2+^ imaging data, we asked whether the increase in presynaptic Ca^2+^ could stem from the inhibition of the Ca^2+^-binding ability of a buffer, resulting in an “un-buffering” of presynaptic Ca^2+^, rather than enhanced Ca^2+^ influx (Extended Data Fig. 2, Extended Data Table 1). Analysis of response kinetics of our experimental GCaMP data revealed significantly faster rise times after baclofen application with no change in decay times (Extended Data Fig. 2A–D). Using GCaMP6s as a model buffer, we simulated Ca^2+^ response kinetics at distinct buffer concentrations and found that high concentrations of a high-affinity Ca^2+^ buffer resulted in significantly prolonged decay times and lower peak fluorescence (Extended Data Fig. 2E–H), ruling out Ca^2+^ un-buffering as the GBR-mediated mechanism. Furthermore, alterations in Ca^2+^ extrusion also failed to reproduce the observed changes in GCaMP fluorescence after baclofen (Extended Data Fig. 2I, J). Finally, we simulated alterations in Ca^2+^ influx and found that a 2.3-fold increase in Ca^2+^ influx most closely reproduced our observed increase in GCaMP peak fluorescence after baclofen (Extended Data Fig. 2K–M).

**Figure 2:**
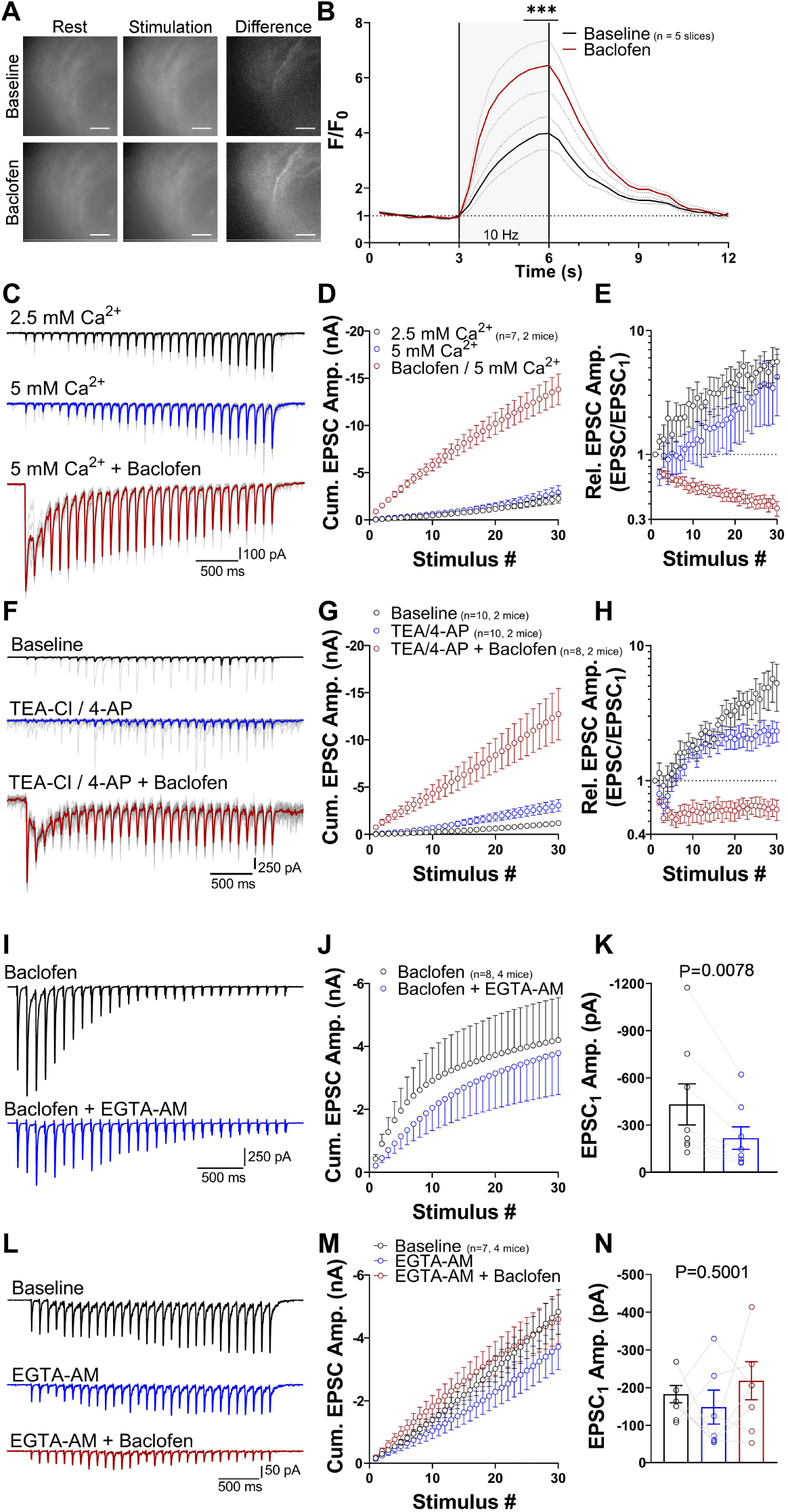
Role of presynaptic Ca^2+^ in tonic and phasic neurotransmission. **A** Example images of axon-GCaMP6s fluorescence at rest (left) and at peak fluorescence during stimulation (middle) under baseline (top) and baclofen conditions (bottom). Subtraction of resting fluorescence from peak fluorescence reveals stimulation-induced fluorescence in a subset of MHb axons (right panels). Scale bars, 50 μm **B** Quantification of GCaMP6s fluorescence time course during stimulation under baseline and baclofen conditions in five slices from five mice. *** P < 0.001 two-way ANOVA with Bonferroni post hoc test **C** Example traces of 10–Hz responses at 2.5 mM and 5 mM Ca^2+^ and at 5 mM Ca^2+^ + 1 μM baclofen. **D** Cumulative EPSC amplitude quantification reveals tonic release at 5 mM Ca^2+^ and a strong transition to phasic release after the addition of baclofen. **E** Normalization of EPSC responses to the first EPSC in the train shows augmentation in 2.5 and 5 mM Ca^2+^ condition and depletion at 5 mM Ca^2+^ + baclofen. **F – G** K^+^ channel blockers TEA and 4-AP did not induce phasic release and did not occlude phasic release induction by baclofen. **I** Example trace of a phasic response before and 10 min after the application of 100 μM EGTA-AM. **J** Cumulative EPSC amplitude plot before and after EGTA-AM application. **K** Comparison before and after EGTA-AM application reveals significant reduction in EPSC1 amplitude. P value calculated with Wilcoxon test. **L – N** Application of EGTA-AM prior to baclofen prevents phasic release induction but does not significantly alter tonic release and its EPSC1 amplitudes. P value calculated by one-way ANOVA. See also Extended Data Fig. 2 and Extended Data Table 1

Since our simulation confirmed increased Ca^2+^ influx as the cause of the GBR-mediated increase in presynaptic GCaMP fluorescence, we next tested whether increasing Ca^2+^ influx by other means was sufficient to induce phasic release. Surprisingly, elevation of extracellular Ca^2+^ from 2.5 to 5 mM or addition of the voltage-gated K^+^ channel blockers tetraethylammonium chloride (TEA-Cl, 1 mM) and 4-aminopyridine (4-AP, 100 μM) neither induced phasic release nor occluded its induction by baclofen (Fig. 2C–H). Although TEA/4-AP application strongly enhanced spontaneous neurotransmitter release, evoked EPSC responses remained tonic and exhibited augmentation (Fig. 2F–H). Thus, we conclude that increasing Ca^2+^ influx alone is insufficient to induce phasic release.

We next hypothesized that increased Ca^2+^ influx during phasic release might result in diffusion of Ca^2+^ further away from the Ca^2+^ channel to recruit SVs more distant from the AZ compared to those for baseline tonic release. Therefore, we tested the effect of a slow Ca^2+^ buffer, EGTA-AM (100 μM), on phasic release, first applied after baclofen (Fig. 2I). EGTA-AM selectively reduced the EPSC amplitudes of early responses in the 10–Hz train but the phasic release pattern remained similar (Fig 2J). Specifically, EGTA-AM reduced the first EPSC (EPSC_1_) amplitude on average by 51.8 ± 2.1% (n=8, 4 mice), suggesting that the phasic release is loosely coupled to Cav2.3^17^. In contrast, EGTA-AM had no significant effect on baseline EPSC1 amplitudes (Fig. 2L-N), suggesting tight coupling for SVs participating in tonic release. Most strikingly, however, EGTA-AM completely blocked the potentiation of EPSC_1_ amplitudes when applied before baclofen (Fig. 2L-N), suggesting that the induction of phasic release strongly depends on Ca^2+^ distant from Cav2.3. Although the exact presynaptic concentration of EGTA is unknown in EGTA-AM experiments, it is unlikely that presynaptic EGTA concentrations differed between tonic and phasic release since no modulation of endogenous esterase activity by GBRs has been known. Therefore, these results support our hypothesis that diffusion of Ca^2+^ away from Ca^2+^ channels recruits loosely-coupled SVs to the RRP of phasic release.

In summary, our results suggest that 1) activation of GBRs on MHb terminals induces an increase in Ca^2+^ influx; 2) this Ca^2+^ increase is necessary for the recruitment of phasic SVs from distal sites of the terminal inducing phasic release; 3) however, increasing Ca^2+^ influx alone is not sufficient to induce phasic release; 4) the tonic SVs are more tightly coupled to Cav2.3 than phasic SVs, indicating distinct release sites for these SVs.

### Comparison of release properties of tonic and phasic neurotransmission

To determine whether GBR-mediated enhancement of release is ascribable to an increase in P_r_, RRP or both, we next compared release properties of basal and GBR-enhanced release by applying 30 stimulations at 100 Hz (Fig. 3A, B). Experiments were performed in the presence of 1 mM kynurenic acid (KA) to avoid AMPA receptor desensitization. The 100–Hz stimulation resulted in the depletion of baseline responses, allowing for the measurement of the tonic RRP in cumulative EPSC amplitude plots^18^. Strikingly, the application of baclofen increased the RRP on average 4.1-fold (Fig. 3C; baseline RRP: 0.58 ± 0.09 nA; baclofen RRP: 2.37 ± 0.36 nA; n=16 recordings/4 mice). In addition, baclofen also increased Pr on average 2.0-fold (Fig. 3D; baseline: 0.086 ± 0.017; baclofen: 0.169 ± 0.024). Altogether, our functional data suggest that tonic and phasic release modes are mediated by two distinct SV pools with different RRP size and coupling tightness, potentially operating in parallel.

**Figure 3:**
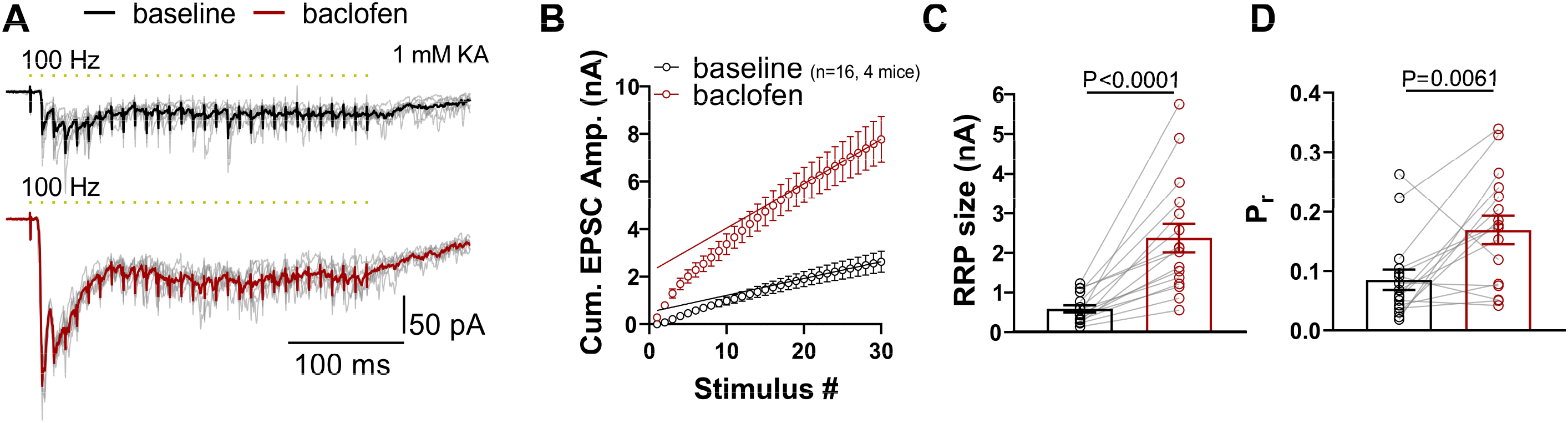
Comparison of RRP and Pr between tonic and phasic release. **A** Example traces of EPSC responses during 100–Hz stimulation at baseline and after baclofen application. **B** Baseline and baclofen cumulative EPSC amplitude plots with linear regression fit through the last 6 stimuli (#25 – #30). **C** Paired comparison of RRP sizes before and after baclofen. P value derived from Wilcoxon t-test. **D** Paired comparison of Pr at baseline and after baclofen, P values calculated by two-tailed unpaired t-test.

### Structural correlates of enhanced neurotransmitter release

Our functional data indicated that phasic release is associated with a 4-fold increase in RRP size compared to basal tonic release (Fig. 3). The RRP is an electrophysiological property considered to be structurally reflected by docked SVs in the AZ^19,20^. To test this, we next used timed high-pressure freezing after optogenetic stimulation of MHb terminals in acute slices (“Flash and Freeze”^21^). First, we measured optogenetic responses in IPN neurons of ChAT-ChR2-EYFP mice^15^ in 200–μm thick coronal slices (Fig. 4A). In ChAT-ChR2-EYFP mice, we found that 90.2% of all asymmetrical synapses expressed channelrhodopsin2 in the rostral/central IPN region (Extended Data Fig. 3A). Under baseline condition, optogenetic stimulation at 10 Hz produced tonic-like EPSC responses that decayed over time after the 10–Hz train (Fig. 4A, B). However, in contrast to electrically-evoked EPSCs, the first baseline EPSC in the optogenetic train was frequently larger than the subsequent responses. This likely resulted from the desensitizing kinetics of channelrhodopsin which, upon activation from resting state, exhibits an initially larger conductance than during steady-state or repetitive activity^22,23^. Importantly, the application of baclofen induced phasic release, which rapidly depleted during the 10–Hz train and recovered within 10 seconds (Fig. 4A, B).

**Figure 4:**
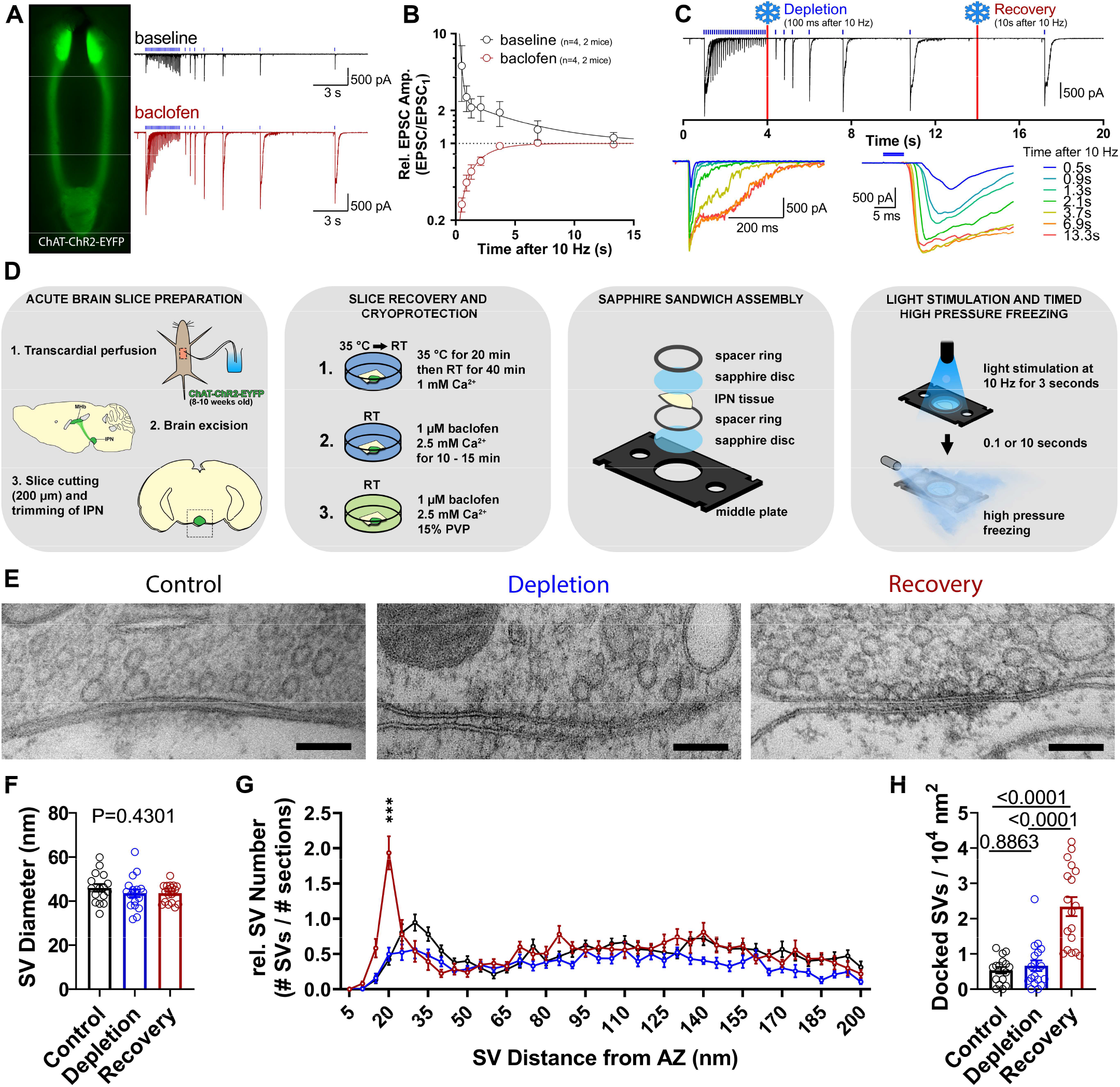
Structural correlates of enhanced neurotransmitter release. **A** EYFP Fluorescence in an angled ChAT-ChR2-EYFP slice (left) and tonic and phasic example traces evoked by optogenetic stimulation in a cell before and during the application of baclofen. **B** Recovery time courses of normalized EPSC amplitudes after augmentation (baseline) and depletion (baclofen). **C** Top, expanded phasic example trace from panel A highlighting freezing time points for depletion and recovery groups in the presence of baclofen. Bottom left, overlay of individual EPSCs over the course of recovery. Bottom right, overlay of individual EPSC onsets during phasic release recovery. **D** Step-by-step scheme displaying the “Flash and Freeze” experiments up to the freezing step. **E** Example EM images of synapses from Control, Depletion and Recovery groups. Scale bars, 100 nm **F** Quantification of synaptic vesicle (SV) diameters across groups. P value derived from one-way ANOVA. **G** Plot of SV numbers (normalized to the number of analyzed serial sections) in bins of 5–nm from the AZ membrane. *** P < 0.001 Recovery vs. Control and Depletion, two-way ANOVA with Bonferroni post hoc test. **H** Quantification of docked SV densities. Values above bars indicate p values calculated by one-way ANOVA with Tukey post hoc test. See also Extended Data Fig. 3

To distinguish tonic and phasic RRPs, we chose to freeze MHb terminals at two time points in the presence of baclofen; 1) 100 ms after the 10–Hz stimulation (Depletion group) at which time point the phasic RRP should be mostly depleted, and 2) 10 s after the 10–Hz stimulation (Recovery group) at which time point the phasic RRP should have fully recovered (Fig. 4C, D). Control samples were prepared in the same way but without light stimulation or baclofen. Following freeze substitution, we extracted the rostral/central IPN region and prepared serial ultrathin sections (40 nm) for EM imaging (Extended Data Fig. 3B–C; Fig. 4E). We measured SV diameters and found no significant difference between groups (Control: 46.0 ± 1.7 nm, n=16 synapses, 3 mice; 2215 SVs measured; Depletion: 43.4 ± 1.7 nm, n=18 synapses, 3 mice, 2204 SVs measured; Recovery: 43.6 ± 0.9 nm, n=18 synapses, 3 mice, 2539 SVs measured; Fig. 4F). In a defined perimeter around the AZ (see Methods), we then measured the distance from each SV’s center to the nearest presynaptic membrane in 5–nm bins. SVs in direct contact with the presynaptic membrane or whose center was within 20–nm distance of the inner leaflet of the membrane were considered docked. In control samples, the peak bins of SVs were in close proximity of the AZ (20 – 40 nm from the membrane) without making contact, thus, considered as loosely-docked^8^ (5^th^ – 8^th^ bins in Fig. 4G). The depletion group displayed lower absolute numbers of SVs, particularly at the loosely-docked bins. In contrast, terminals in the recovery group showed a strong increase in docked SVs (first four bins in Fig. 4G). Furthermore, the density of docked SVs in the recovery group (2.34 ± 0.27 docked SVs / 10^4^ nm^2^; n=18 terminals, 3 mice) increased 3.5-fold compared to the depletion group (0.67 ± 0.15 docked SVs / 10^4^ nm^2^; n=18 terminals, 3 mice; Fig. 4H). The difference in docked SV densities between depletion and recovery groups was similar to the difference in RRP size between tonic and phasic release (4.1-fold, Fig. 3C), supporting that two distinct pools of SVs mainly contribute to each release mode.

### SV-associated molecules selectively involved in tonic and phasic release

To test the two-pool mechanism hypothesis, we next aimed to identify vesicle-associated proteins selectively involved in tonic and phasic release. We first investigated the role of synaptoporin (SPO) based on its selective expression in ventral MHb neurons (Allen Brain Atlas). SPO is a vesicular membrane protein and a homologue of synaptophysin^24^, but its functional importance remains unknown^25^. In contrast to synaptophysin, SPO was found to co-precipitate with Cav2.3 in a brain-wide proteomics study^26^, suggesting that it may be relevant for neurotransmission in MHb terminals which exclusively rely on Cav2.3 for release^1^. Immunofluorescence revealed that SPO was expressed in axon-like structures in the rostral/central IPN subnuclei (Fig. 5A), and at the EM level, SPO-labeling was confirmed on SVs in the vast majority of MHb terminals (Fig. 5B, Extended Data Fig. 4A). We then generated SPO KO mice (see Method) and confirmed the specificity of immunolabeling (Fig. 5A). To test the functional role of SPO, we performed whole-cell recordings from IPN neurons in SPO KO mice (Fig. 5C–F). At baseline conditions, augmentation of tonic release appeared strongly impaired compared to wild-type (WT) mice (Fig. 5D). In contrast, baclofen-induced phasic release did not appear different from that of WT and P_r_ of EPSC_1_ remained unaltered (Fig. 5E, F; measured by linear regression analysis of cumulative EPSC amplitude plots). Although there was a significant effect of genotype on phasic Pr time course (F_1, 690_ = 7.599, P = 0.006, two-way ANOVA), Bonferroni post hoc test revealed no significant difference at any stimulation time point. These results suggest that SPO is a mediator of tonic release augmentation and thus, could be a molecular marker specific for tonic release SVs.

**Figure 5:**
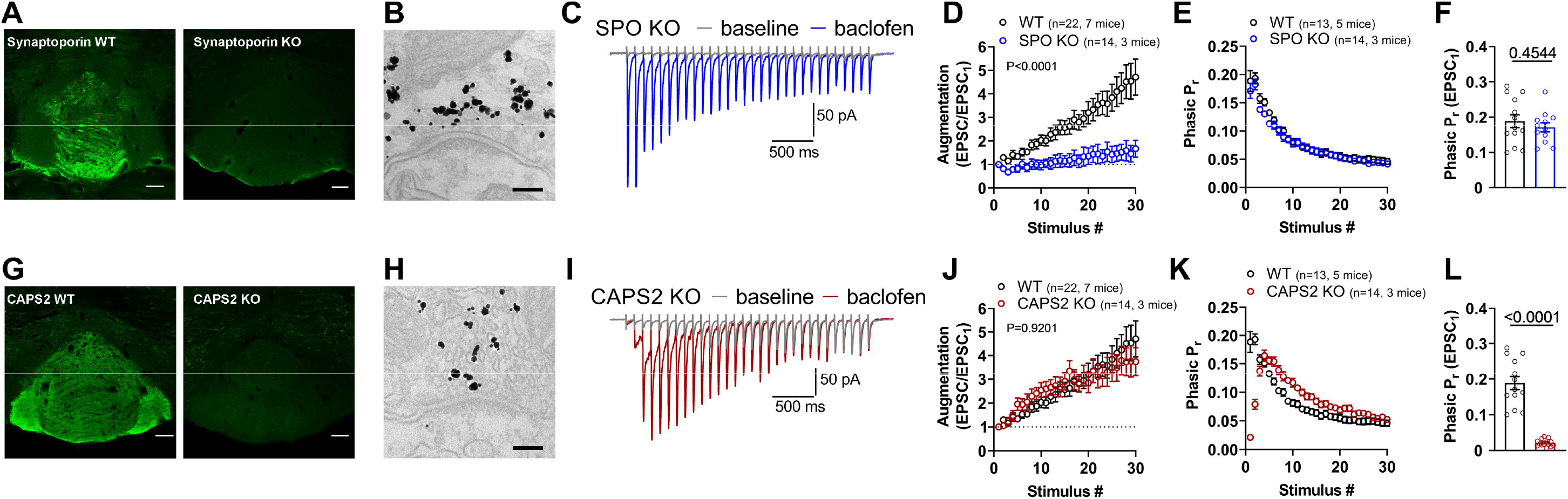
SV-associated molecules involved in tonic and phasic neurotransmitter release. **A** Confocal images of anti-synaptoporin immunofluorescence labeling in the IPN of a WT and a SPO KO mouse. Scale bars, 100 μm. **B** Pre-embedding immunolabeling for SPO in an MHb terminal, showing gold particles co-localized with SVs. Scale bar, 100 nm. **C** Example 10–Hz EPSC traces of a recording in an acute slice from a SPO KO mouse at baseline and during baclofen application. **D** Tonic release augmentation was significantly impaired in SPO KO mice, displayed P value of main effect of genotype, calculated by two-way ANOVA. Bonferroni post hoc analysis revealed a significant difference in stimuli #20 – #30. **E** Overlay of phasic release Pr time course between WT and SPO KO mice, calculated by linear regression analysis of cumulative EPSC amplitude plots^18^. **F** Pr of EPSC1 in the phasic response train was not different between WT and SPO KO mice. P value calculated from two-tailed unpaired t-test. **G** Confocal images of anti-CAPS2 immunofluorescence labeling in the IPN of a WT and CAPS2 KO mouse. Scale bars, 100 μm **H** Pre-embedding immunolabeling for CAPS2 in a MHb terminal shows gold particles co-localized with SVs. Scale bar, 100 nm. **I** Example 10 Hz EPSC traces recorded in an acute slice from a CAPS2 KO mouse at baseline and during baclofen application. **J** Augmentation remained unaffected by genetic ablation of CAPS2. Main effect of genotype P value calculated via two-way ANOVA **K** Pr time course of phasic release, calculated by linear regression analysis of cumulative EPSC amplitude plots, was strongly affected by genetic ablation of CAPS2 compared to WT. **L** Pr of EPSC_1_ in the phasic response was significantly lower in CAPS2 KO compared to WT mice. P value calculated by two-tailed unpaired t-test. See also Extended Data Fig. 4 and Extended Data Fig. 5

Phasic release requires the cytosolic Ca^2+^-dependent activator protein for secretion (CAPS) in the hippocampus^8,27^. Although there are two CAPS isoforms, CAPS1 and CAPS2, only CAPS1 is required for fast, phasic neurotransmission in the hippocampus^27^, whereas CAPS2 was found to be involved in the release of neuropeptides and neurotrophic factors^28,29^. Since MHb neurons exclusively express CAPS2 but not CAPS1^30^, we hypothesized that CAPS2 might be involved in GBR-induced phasic release. Immunofluorescence revealed strong CAPS2 expression in all IPN subnuclei (Fig. 5G) and pre-embedding EM showed CAPS2 labeling on SVs in the vast majority of MHb terminals in the rostral/central IPN (Fig. 5H, Extended Data Fig. 4B). Double immunofluorescence showed a large overlap of labeling for CAPS2 with that for SPO (Extended Data Fig. 4C). We then performed whole-cell recordings in CAPS2 KO^27^ (Fig. 5I–L). Functionally, tonic release appeared intact, with augmentation similar to that of WT (Fig. 5I, J). However, phasic release was strikingly altered in CAPS2 KO mice (Fig. 5K, L). Specifically, Pr of EPSC1 was reduced almost 10-fold compared to WT (WT EPSC1 Pr: 0.19 ± 0.02, n=13, 5 mice; CAPS2 KO EPSC1 Pr: 0.02 ± 0.00, n=14, 3 mice, Fig. 5L). Furthermore, P_r_ increased over the first four stimuli before starting to reduce gradually. Interestingly, we observed similar phasic responses in WT mice but only at the very first 10–Hz stimulation in the presence of baclofen (Extended Data Fig. 5A, B). Therefore, these results suggest that CAPS2 is recruited Ca^2+^-dependently during phasic release induction and might serve to stabilize replenished SVs in a docked state. Nonetheless, the facilitation in the first several stimuli followed by depression in CAPS2 KO mice (Fig. 5K, L) indicates that CAPS2 does not interfere with the Ca^2+^-dependent recruitment of phasic release SVs. Overall, we identified SPO and CAPS2 as SV-associated proteins selectively involved in tonic and phasic release, respectively.

### Activity-dependent induction and retention of phasic release

Our results in CAPS2 KO mice suggest that CAPS2 is required for the retention of newly recruited SVs in the RRP, but previous activity is necessary for the potentiation of initial EPSCs in the train. Therefore, we tested whether achieving the maximal phasic response is activity-dependent or whether baclofen without activity is sufficient to reach peak potentiation. To this aim, we recorded bilateral MHb inputs to single IPN neurons and, after establishing a baseline response, stopped stimulation on one hemisphere while washing in baclofen (silent washin, Fig. 6A, B). Once the 10–Hz stimulation (3 s, every 20 s) on the other hemisphere induced maximal potentiation, the same 10–Hz stimulation of the silent side resumed (Fig. 6B). Strikingly, EPSC_1_ amplitude in the first 10–Hz stimulation train was only marginally increased whereas the second train exhibited maximal potentiation of EPSC_1_ amplitude. This result confirmed that the induction of phasic release is activity-dependent.

**Figure 6:**
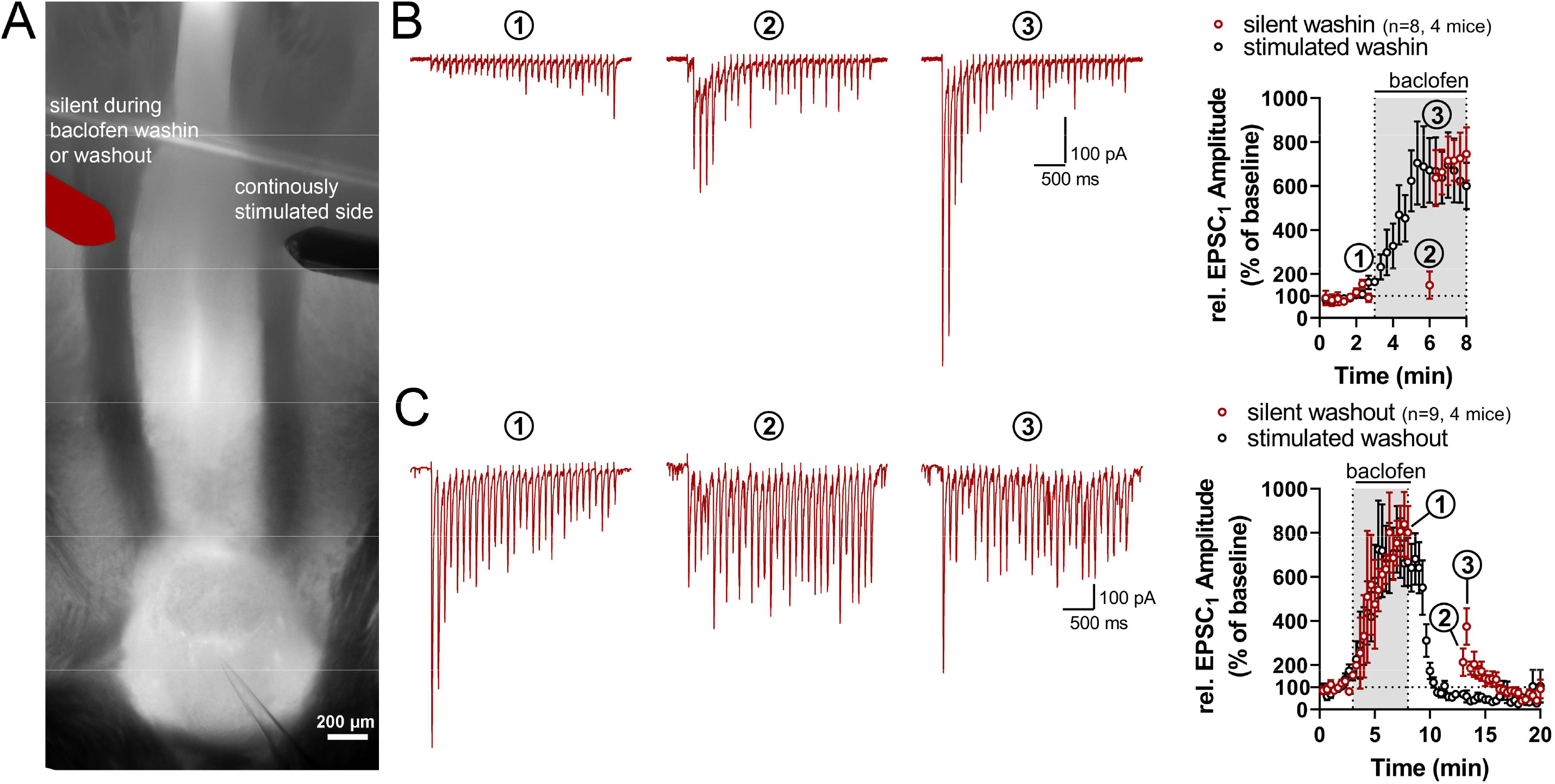
Potentiation by GBR activation and its storage is activity-dependent. **A** Example thick-slice configuration of recording in IPN with bilateral MHb input stimulation. To examine activity-dependence (**B**), a single IPN neuron was patched and after bilateral baseline recordings (10–Hz stimulation for 3 s, 5–s interval between left and right side stimulations, 20 s between sweeps), stimulation with the red-highlighted electrode was halted during baclofen washin, whereas stimulation was continued with the other electrode (10 Hz, 3–s duration, every 20 s). Once maximal potentiation was reached, stimulation with the red-highlighted electrode resumed. To examine storage of potentiation (**C**), after bilateral baseline recording and bilateral induction of maximal potentiation, baclofen washout commenced and stimulation with the red-highlighted electrode was halted. Stimulation with the other electrode continued and, once potentiation was fully reverted, stimulation with the red-highlighted electrode resumed. **B** Left, example traces derived from the silent side at baseline (1), at the first (2) and second (3) 10–Hz stimulus after silent baclofen washin. Right, time course of EPSC_1_ amplitudes, relative to baseline, during stimulated and silent washin of baclofen. **C** Left, example traces derived from the silent side at peak potentiation (1), at the first (2) and second (3) stimulus after silent baclofen washout. Right, time course of relative EPSC_1_ amplitudes during stimulated and silent washout of baclofen.

Next, we asked whether the GBR-mediated potentiation, once induced, might be stored in the absence of GBR activation during periods of synaptic inactivity. To test this, we stimulated MHb inputs bilaterally to induce phasic release on both sides in the presence of baclofen. Once full potentiation was achieved, stimulation on one hemisphere was halted (silent washout) while starting the washout of baclofen (Fig. 6C). Five minutes after the washout onset, potentiation on the continuously stimulated side was fully reverted (stimulated washout). At this time point, we restarted the 10–Hz stimulation of the silent side and saw only marginal potentiation of EPSC_1_ amplitudes in the first train (EPSC1: 213.2 ± 62.3% of baseline amplitude, n=9, 4 mice; Fig. 6C). However, in the second train, EPSC_1_ amplitudes increased to 46.8% of the peak potentiation (second train EPSC_1_: 374.7 ± 82.8% of baseline amplitude; peak potentiation EPSC_1_: 801.6 ± 120.4% of baseline amplitude). Therefore, we conclude that the GBR-mediated potentiation can be stored for minutes in the absence of GBR activation, but the retrieval of this stored potentiation also requires activity, consistent with the idea of Ca^2+^-dependent recruitment of phasic release SVs.

### Recruitment of CAPS2 to the presynaptic active zone by phasic but not tonic release

Given the partial storage of GBR-mediated potentiation for at least five minutes (Fig. 6C), we hypothesized that chemical fixation might be sufficiently fast to capture molecular changes underlying the phasic release induction. Therefore, we next investigated the nano-anatomical location of CAPS2 inside MHb terminals during phasic release. We combined optogenetic stimulation of MHb terminals in acute IPN slices from ChAT-ChR2-EYFP mice and immersion chemical fixation followed by pre-embedding immunolabeling (“Flash and Fix”). Control slices remained unstimulated and unexposed to baclofen whereas the “10 Hz” group received three 10–Hz light stimulations (3 s duration, 10 s intervals) without baclofen and the “10 Hz + Baclofen” group received the same stimulation after a 10 – 15 min pre-incubation in 1 μM baclofen. Immediately after the last stimulation, slices were submerged in a fixative (4% PFA, 0.05% glutaraldehyde, 15% picric acid) for 45 min. Pre-embedding immunolabeling for CAPS2 revealed that CAPS2 was specifically recruited to the presynaptic AZ in the 10 Hz + Baclofen group (Fig. 7A, B; Control: 0.3 ± 0.1 particles/AZ, n=58 profiles, 2 mice; 10 Hz: 0.7 ± 0.1 particles/AZ, n=68 profiles, 2 mice; 10 Hz + Baclofen: 2.6 ± 0.3 particles/AZ, n=60 profiles, 2 mice). This result suggests the recruitment and subsequent stabilization of CAPS2 in the presynaptic AZ during the phasic, but not tonic, release.

**Figure 7:**
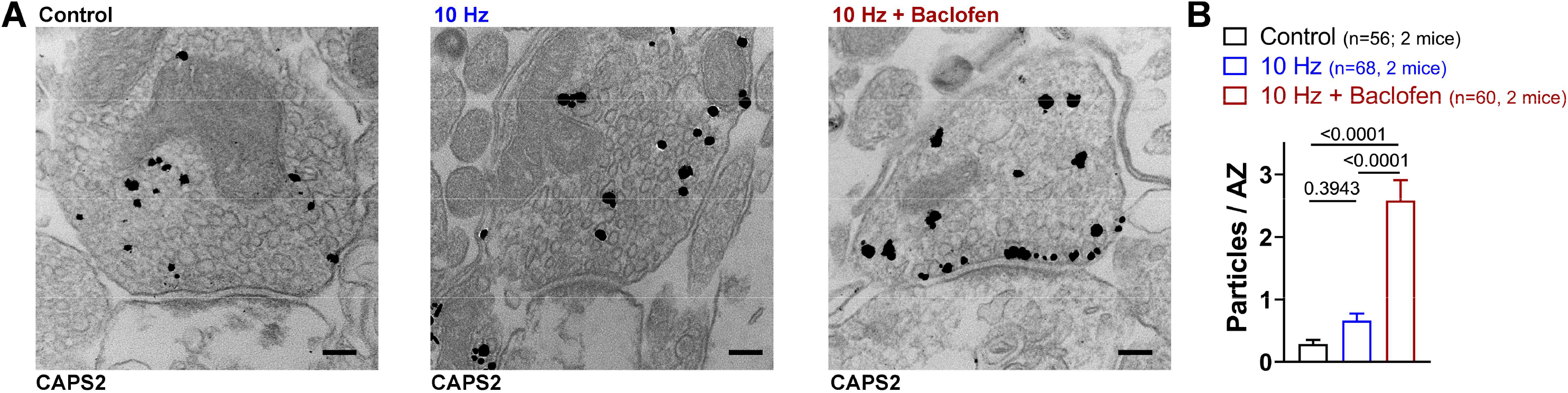
Recruitment of CAPS2 to the active zone during phasic release. **A** Example images of pre-embedding EM labeling for CAPS2 in acute slices under control conditions without stimulation and after 10–Hz stimulation in the absence (10 Hz) or presence of 1 μM baclofen (10 Hz + Baclofen). **B** Quantification of particle numbers in the AZ reveals significantly larger CAPS2 particle numbers in the “10 Hz + Baclofen” group compared to the other groups. P values calculated by one-way ANOVA with Tukey post hoc test.

### “Flash and Freeze-fracture” reveals selective recruitment of CAPS2 to the presynaptic membrane during phasic release

SVs containing SPO or bound by CAPS2 may selectively fuse with the AZ membrane during tonic and phasic release, respectively. However, it is unclear whether all SVs contain SPO and CAPS2 or whether tonic and phasic pools are molecularly non-overlapping. To answer this question, we combined “Flash and Freeze” with freeze-fracture replication and subsequent immunolabeling. This method, which we call “Flash and Freeze-fracture”, enables the nanoscale detection of multiple proteins simultaneously in the presynaptic membrane within milliseconds of neurotransmitter release (Fig. 8A).

**Figure 8:**
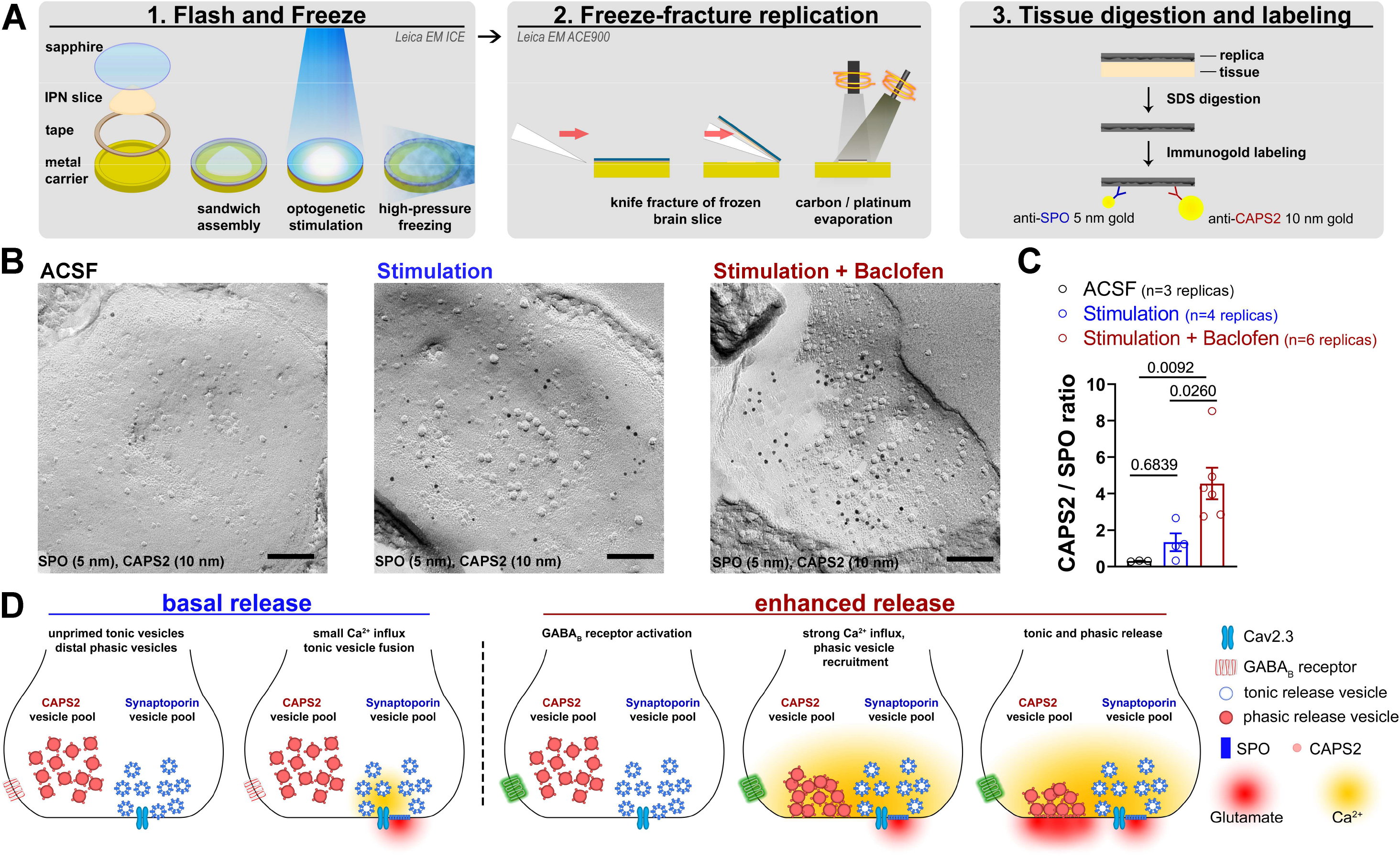
“Flash and Freeze-fracture” replica immunolabeling for SPO and CAPS2 in the presynaptic active zone during tonic and phasic release. A “Flash and Freeze-fracture” steps schematic. Left Panel, “Flash and Freeze” of acute brain slices using sapphire/metal hybrid sandwich separated by a double-sided tape. Middle panel, Freeze-fracture replication of frozen brain slices using a knife to separate sapphire from the metal carrier. Right Panel, SDS treatment of tissue and immunolabeling with primary antibodies against SPO and CAPS2 followed by labeling with 5 nm and 10 nm gold-conjugated secondary antibodies. **B** Example images of SPO and CAPS2 labeling in “Flash and Freeze-fracture” replicas frozen without light stimulation (ACSF) or after an 8 ms light pulse in the absence (Stimulation) or presence of 1 μM baclofen (Stimulation + Baclofen). Scale bars, 100 nm **C** Quantification of CAPS2/SPO particle ratio revealed a significant increase in the “Stimulation + Baclofen” group. P values derived from one-way ANOVA with Tukey post hoc test. **D** Summary scheme depicting a hypothetical two-pool mechanism underlying the transition from the tonic to phasic release mode in MHb terminals. Left: In the basal release condition, tonic SPO-positive SVs are loosely docked in close vicinity of Cav2.3, whereas CAPS2-positive phasic SVs are located more distally and undocked. Action potential-triggered Ca^2+^ influx is small and only closely located SPO-positive tonic SVs fuse with the AZ membrane. Right: In the enhanced release condition, GBR activation coupled with action potential-mediated depolarization produces a massive influx of Ca^2+^, which initially triggers release of closely located SPO-positive tonic SVs but also rapidly recruits larger number of distal CAPS2-positive phasic SVs (middle) to the AZ. The subsequent action potentials trigger the release of both closely located tonic and newly recruited phasic SVs, resulting in an increased RRP (right). Following the fusion of CAPS2-positive phasic SVs, CAPS2 remains membrane-bound and may act as a docking site for replenished phasic SVs, resulting in retention of the increased RRP. See also Extended Data Fig. 6

If CAPS2 is exclusively expressed on phasic and SPO exclusively on tonic SVs, then the ratio of CAPS2 to SPO molecules in the presynaptic membrane should shift towards CAPS2 during phasic release. In contrast, if all SVs contain CAPS2 and SPO, the ratio of CAPS2 to SPO molecules in the presynaptic membrane should remain unaltered between tonic and phasic release. Thus, we next quantified the ratio of CAPS2 to SPO in the presynaptic membrane following stimulation by a single 8 ms light pulse in the absence or presence of baclofen. Since phasic SV recruitment is activity-dependent (Extended Data Fig. 5A and Fig. 6B), two 10–Hz conditioning stimuli (3–s duration, 10–s interval) were given 10 s prior to the single light pulse. Freezing was timed so that the tissue reached 0 °C 8 ms after light onset and light application continued for an additional 15 ms during freezing. To maximize release and stabilize SV- and membrane-associated proteins in the presynaptic membrane after fusion, slices were stimulated and frozen in the presence of 1 mM TEA-Cl, 100 μM 4-AP and 100 μM dynasore. Because CAPS2 is a cytosolic protein, we incubated replicas with attached tissue in 2% PFA for one hour prior to SDS digestion to crosslink CAPS2 to potential interacting transmembrane proteins. In “Flash and Freeze-fracture” replicas of “stimulation + baclofen” group, the ratio of CAPS2 to SPO (4.56 ± 0.87, n=6 replicas, 205 profiles, 6 mice) was significantly higher than in the unstimulated control group (ACSF, 0.31 ± 0.01, n=3 replicas, 153 profiles, 3 mice) and “stimulation” group (1.33 ± 0.48, n=4 replicas, 320 profiles, 4 mice; Fig. 8B, C). We confirmed the specificity of replica labeling for SPO and CAPS2 in acute-slice replicas of corresponding KO mice (Extended Data Fig. 6A). The difference in the ratio during phasic release derived from increases in the density of CAPS2 in the presynaptic membrane (Extended Data Fig. 6B) as SPO density was similar across all three groups (Extended Data Fig. 6B), potentially due to high spontaneous neurotransmission in the presence of TEA/4AP (Fig. 2F). Comparison of SPO and CAPS2 particle numbers in individual AZs revealed significant positive correlation in the “stimulation + baclofen” group but not the “stimulation” group (Extended Data Fig. 6C), indicating that baclofen induced a correlated evoked vesicular release of SPO- and CAPS2-associated SVs, whereas release of these two populations was independent in the stimulation only condition. Although control samples also showed a significant correlation, this was likely caused by the frequent absence of CAPS2 in many of the control profiles. Finally, to test for association of SPO and CAPS2 in the AZ, we compared the nearest neighbor distances (NNDs) from CAPS2 to SPO particles with those from real CAPS2 to simulated SPO particles (see Methods). Interestingly, NND analysis revealed no significant association of CAPS2 and SPO in the presynaptic membrane of both “stimulation” and “stimulation + baclofen” groups (Extended Data Fig. 6D), suggesting that fusion sites of CAPS2- and SPO-associated vesicles may not overlap.

## Discussion

We discovered that activation of GBRs induces a transition from tonic to phasic neurotransmitter release mode in MHb terminals, which coincides with robust potentiation and an increase in action potential-driven Ca^2+^ influx as previously reported^3,4^. During basal release, small Ca^2+^ influx triggers release of tonic SVs close to the VGCC without involving the distal phasic pool. Importantly, we found GBR-mediated recruitment of additional SVs from distal presynaptic regions, resulting in increases in RRP size and docked SVs. We identified SPO and CAPS2 as two SV-associated molecules selectively involved in tonic and phasic release, respectively. The recruitment of phasic release SVs is activity-dependent and their retention at the AZ is CAPS2-dependent. Strikingly, a newly developed “Flash and Freeze-fracture” method revealed selective transfer of CAPS2 to the presynaptic membrane during phasic release. Thereby, we propose a novel two-pool mechanism (Fig. 8D) underlying the unusual GBR-mediated potentiation of neurotransmitter release from MHb terminals.

To our knowledge, all currently known synapses release exclusively in a tonic or phasic manner at a given frequency^8,9^. The MHb terminal in the IPN is the only example capable of a rapid and reversible transition between tonic and phasic release modes at the same release frequency. The combination of our functional and structural data indicates that the two modes co-exist in the same terminals with two distinct populations of SVs selectively involved in these two modes. First, the Ca^2+^ imaging of MHb terminals (Fig. 2A-B) showed increased Ca^2+^ signals in all responsive axons but detected no new axons appearing after GBR activation, indicating that the same axons are involved in both tonic and phasic release. Second, “Flash & Freeze” revealed an increase in docked SVs in the vast majority of terminals after recovery from phasic release depletion, whereas the docked SV numbers were similarly low in the control and depletion conditions (Fig. 4H), indicating that single population of terminals increase docked SVs in phasic mode and depleted them to the same level as tonic mode. Third, “Flash & Freeze-fracture” showed that the increased CAPS2 labeling in the phasic mode (Stimulation + Baclofen) always co-localized with SPO labeling at the AZ (Fig. 8B-C, Extended Data Fig. 6C), indicating that the same terminals have CAPS2- and SPO-associated SVs. Fourth, SPO- and CAPS2-deficient mice have selective impairments in tonic and phasic release, respectively (Fig. 5), but the vast majority of MHb terminals were labeled for SPO and CAPS2 with a large overlap (Extended Data Fig. 4), indicating that tonic and phasic SVs co-exist in the same terminals. Fifth, tonic and phasic SVs have distinct coupling tightness, indicating different distances from Cav2.3 in the AZ. Sixth, “Flash & Freeze-fracture” showed a constant level of SPO at the AZ in the presence and absence of baclofen (Extended Data Fig. 6B), indicating GBR-independent release of SPO-associated SVs in both tonic and phasic modes, whereas the drastic increase in CAPS2 number (Fig. 7) and CAPS2/SPO ratio (Fig. 8B-C) by baclofen indicates selective recruitment of CAPS2-associated SVs in the phasic mode. In addition, the positive correlation between SPO and CAPS2 numbers in each AZ in the presence but not the absence of baclofen (Extended Data Fig. 6C) suggests that evoked co-release of SPO- and CAPS2-associated SVs occurs only in phasic release. No significant co-localization of CAPS2 and SPO in the AZ membrane also supports separated fusion sites for the two SV populations. These results provide strong support for the two-pool mechanism with distinct SV populations, SPO- and CAPS2-associated ones mediating tonic and phasic release, respectively, in the same MHb terminals.

We interpret the convergence of tonic and phasic release to similar EPSC amplitudes (Fig. 1D) as an indicator of their parallel nature. However, this assumes that the phasic pool depletes completely, which is not described in other phasic synapses at low stimulation frequencies. On one hand, lack of complete depletion in other synapses might be caused by the presence of an “invisible” tonic component that is masked by the depressing phasic pool^16^. On the other hand, in addition to a parallel SV pool model, our results could also be interpreted according to a recently proposed two-step sequential docking model which categorizes SVs into different docking categories but draws them from a single, molecularly homogenous pool^8,31,32^. Under baseline conditions, the majority of SVs may be in a loosely docked state (SV_LS_) with only a small proportion SVs in a tightly docked, fusion-competent configuration (SV_TS_). With repeated stimulation, SVs may gradually transition Ca^2+^-dependently from LS to TS to fusion until a steady-state is reached. Accordingly, baclofen may shift the resting ratio of LS/TS pools towards TS, resulting in a higher proportion of tightly docked SVs at rest. Although the starting pool ratios of LS/TS in tonic and phasic release are different, repeated stimulation in the presence of baclofen would still lead to the same steady-state as during tonic release, resulting in apparent conversion of facilitated tonic and depressed phasic EPSC amplitudes. According to the sequential model, the release pattern of the loosely-docked pool closely matches our partial phasic responses to the first repetitive stimulation in WT and those to all repetitive stimulations in CAPS2 KO. This could suggest that 1) SV transition from a loosely to a tightly docked state during phasic release induction and 2) CAPS2 mediates the tight docking required for the full phasic release. Importantly, however, a selective recruitment of CAPS2 to the AZ during phasic but not tonic release is difficult to explain with a molecularly homogenous SV pool from which all SVs are drawn. The distinct coupling distance of tonic and phasic SVs and no significant co-localization of SPO and CAPS2 in the AZ also favor the parallel model.

Unexpectedly, we identified SPO as a mediator of tonic release augmentation. Until now, no clear functional role of SPO has been described^25^. Synaptoporin (also called synaptophysin 2) is a homologue of synaptophysin with several molecular distinctions, one of which is the direct binding to Cav2.3 R-type Ca^2+^ channels^26^. It is conceivable that the interaction of SPO with Cav2.3 in MHb terminals underlies the tighter coupling of tonic compared to phasic release vesicles. Furthermore, Cav2.3 might serve as an anchor point for tonic vesicles during augmentation and loss of this anchor might result in impaired recruitment of tonic vesicles to the AZ during tonic release.

Similar to SPO, the function of the cytosolic CAPS2 protein in fast neurotransmission has remained elusive. Although the more widely expressed CAPS1 is a known mediator of phasic release in the hippocampus, CAPS2 KO alone did not induce any alterations in fast neurotransmission^27^. Instead, CAPS2 was found to serve as a mediator of neuropeptide and neurotrophic factor release during development in other brain regions^30,33’^. Therefore, we cannot rule out developmental alterations in MHb terminals of CAPS2 KO. The CAPS2 function found in MHb terminals, which exclusively express CAPS2 isoform^30^, may be masked in other synapses expressing both CAPS1 and CAPS2. Approximately 40% of CAPS proteins elude in the membrane fraction in whole-brain lysates and synaptosomal preparations^34^, suggesting that they are often membrane-bound. Importantly, CAPS2 is also located on SVs in parallel fiber terminals in the cerebellum^33^. Thus, the recruitment of CAPS2 to the AZ we observed during phasic activity could result from the fusion of CAPS2-associated SVs, although we cannot rule out that cytosolic CAPS2 is recruited to the AZ by other mechanisms. Most strikingly, in WT MHb terminals, phasic release induction was activity-dependent, with the first repetitive stimulation showing a partial phasic response (Extended Data Fig. 5A) but inducing a fully potentiated response to the subsequent repetitive stimulation. In contrast, in CAPS2 KO mice, the repetitive stimulation induced the partial phasic response every time as if it were the first stimulation. These results suggest that phasic vesicles are activity-dependently recruited to the AZ and, upon fusion with the membrane, CAPS2 proteins associated with the fused vesicles remain at the fusion site, potentially via the Ca^2+^ binding C2 and the plexstrin homology (PH) domains^35^. The remaining CAPS2 at the AZ might serve as docking sites for replenished vesicles via their Munc13 homology (MHD) and MUN domains^35^, resulting in an increase in the RRP. Contrary to this hypothesis, a previous study in CAPS1/2-deficient hippocampal cultured neurons found that the PH but not the MUN domain was required for CAPS2-mediated RRP recruitment^36^. In the CAPS2 KO, phasic vesicles have to be recruited to the AZ “from scratch” at every stimulus train, which may be interpreted as synaptic short-term memory loss of prior activity and thus, CAPS2 might be viewed as an engram molecule^37^.

Our results suggest that the same somatic firing pattern in the MHb *in vivo* may produce two distinct output signals from IPN neurons. Specifically, phasic release likely induces short-duration burst firing in IPN neurons followed by periods of silence, whereas tonic release may produce prolonged, uninterrupted firing. A main IPN projection target is the median raphe nucleus (MRN)^38–40^, a serotonergic center that modulates a multitude of forebrain functions^41^. Interestingly, tonic excitation of MRN neurons via the IPN is hypothesized to disrupt hippocampal theta oscillation through MRN-derived serotonergic innervation^42^. In addition, high-frequency firing GABAergic MRN neurons have been shown to be active in either tonic or phasic rhythms^43^. Similarly, the IPN also projects to the lateral habenula neurons^40^ which exhibit a tonic excitation and phasic inhibition patterns during motivation-related behaviors^44^. Interestingly, downregulation of CAPS2 in MHb neurons induces depression-like phenotypes in mice^45^. Thus, tonic and phasic IPN output patterns might modulate forebrain serotonergic signaling through control of raphe activity as well as motivational states through lateral habenula projections.

Furthermore, activation of GBRs on MHb terminals might selectively strengthen inputs from low-frequency (1 Hz) firing cells, as terminals of high frequency (10 Hz) firing MHb cells might rapidly revert to basal release strength following RRP depletion. What could be the source of GABA for the transition to phasic release? Since the vast majority of IPN neurons are GABAergic and innervate each other together with MHb-derived cholinergic inputs^46,47^, local GABA release from presynaptic or postsynaptic IPN neurons^3^ might regulate their MHb-driven output through the activation of presynaptic GBR_S_. Although the trigger for retrograde GABA release is unknown, a coincidence detection mechanism may be envisioned where secondary excitatory projections to the IPN^48^ raise ambient GABA concentrations high enough to activate GBRs on MHb terminals.

The surprising two-pool mechanism we discovered raises several questions: Which factors retain CAPS2 in the AZ during phasic release? What is the exact reason for the increased influx of Ca^2+^ during GBR activation? What is the GBR-mediated signaling mechanism necessary for the phasic release induction other than the Ca^2+^ increase? Future studies will be necessary to answer these questions. Overall, our study provides new insights into the mechanisms underlying the unusual potentiation of release by GBRs from MHb terminals and opens up a new regime of presynaptic modulation.

## Materials and Methods

### Animals

Wild-type C57BL6J (#000664), heterozygous ChAT-ChR2-EYFP (#014546) and heterozygous ChAT-IRES-Cre (#006410) mice were purchased from Jackson Laboratory. Homozygous CAPS2 KO mice were generated as previously described^27^. Synaptoporin (MGI:1919253) KO mice were generated using the CRISPR-Cas9 method with AAGTACCGCGAAAACAACCGGGG as a guide RNA target sequence. The single-guide RNA and Cas9 protein were co-injected into fertilized oocytes, collected from superovulated C57BL/6J female mice mated with male mice. All the surviving zygotes were transferred into oviductal ampullae of pseudopregnant recipient mice. To determine the genotype of pups, genomic DNA was extracted from ear tissues, and screened by PCR analysis and subsequent direct sequencing analyses. A homozygous 2-bp deletion at amino acid position 130 resulted in a STOP codon at amino acid position 136, which corresponds to the start of the third transmembrane domain^49^. Male and female mice were used indiscriminately in all experiments. Mice were bred and maintained at the preclinical facility at Institute of Science and Technology Austria on a 12h light/dark cycle with access to food and water *ad libitum.* All experiments were performed in strict accordance with the license approved by the Austrian Federal Ministry of Science and Research (Animal license number: BMWFW-66.018/0012-WF/V/3b/2016) and Austrian and EU animal laws.

### Electrophysiology

Mice (6 – 12 weeks of age) were deeply anesthetized via intraperitoneal (i.p.) injection of ketamine (90 mg/kg) and xylazine (4.5 mg/kg), followed by transcardial perfusion with ice-cold, oxygenated (95% O_2_, 5% CO_2_) artificial cerebrospinal fluid (ACSF) containing (in mM): 118 NaCl, 2.5 KCl, 1.25 NaH_2_PO_4_, 1.5 MgSO_4_, 1 CaCl_2_, 10 Glucose, 3 Myo-inositol, 30 Sucrose, 30 NaHCO_3_; pH = 7.4. As described previously ^1^, the brain was rapidly excised and a single, 52–54° angled slice of 1 mm thickness containing the whole MHb to IPN pathway was cut using a Pro7 Linear Slicer (Dosaka, Kyoto, Japan). Slices were left to recover for 20 min at 35°C, followed by a slow cool down to room temperature (RT, 22.5 – 24.0 °C) over 40 – 60 min. After recovery, the slice was transferred to the recording chamber and superfused with ACSF containing 2.5 mM CaCl_2_ and 20 μM bicuculline methiodide (Tocris, Bristol, UK) at a rate of 3 – 4 ml/min at RT. Glass pipettes (B150-86-10, Sutter Instrument, Novato, CA, USA) with resistances of 3 – 4 MΩ were crafted using a P1000 horizontal pipette puller (Sutter Instrument) and filled with internal solution containing (in mM): 130 K-Gluconate, 10 KCl, 2 MgCl2, 2 MgATP, 0.2 NaGTP, 0.5 EGTA, 10 HEPES, 5 QX314-Cl; pH 7.4 adjusted with KOH. Neurons of the rostral and central IPN were visually identified using an infrared differential interference contrast video system in a BX51 microscope (Olympus, Tokyo, Japan). Electrical signals were acquired at 20 – 50 kHz and filtered at 4 kHz using a Multiclamp 700B amplifier connected to a Digidata 1440A digitizer (Molecular Devices, San Jose, CA, USA) with pClamp10 software (Molecular Devices). Stimulating electrodes (CBBPC75, FHC, Bowdoin, ME) were placed bilaterally on the fasciculus retroflexus and electrical stimulation (10 Hz, 3 seconds, 0.2 ms pulse duration, 0.5 – 2.5 V stimulation intensity) of left or right habenular axon fiber tracts was applied via a stimulus isolator (AMPI, Jerusalem, Israel). Neurons were voltage-clamped at –60 mV in whole-cell mode. Recordings with access resistances exceeding 20 MΩ or with changes in access resistance or holding current by more than 20% were discarded. Access resistance was not compensated. At each sweep, left and right fiber tracts were stimulated separately with 5–s intervals and inter-sweep intervals of 20 s. Left- and right-side derived EPSC responses were treated as individual recordings, meaning that one whole-cell recording of one IPN neuron receiving two independently stimulated inputs would yield n = 2. Tonic and phasic release traces were analyzed after averaging the responses of 5 – 10 sweeps. EPSC amplitudes were measured as the maximal negative current deflection between two stimulation artifacts (or within 100 ms in case of the last stimulus). For time course measurements of recovery from augmentation and depletion, the left or right fasciculus retroflexus was stimulated in separate recordings at 10 Hz for 3 s followed by a single stimulus at intervals from 0.5 – 13.3 s. To calculate RRP size and P_r_ using cumulative EPSC amplitudes, a linear correlation was fit through the last six points of the cumulative EPSC amplitude plot (stimulus #25 – stimulus #30) and the projected value of the line at x=1 was considered the RRP size (I_RRP_). Release probability was calculated as 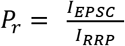. At 100–Hz trains, the latency of the first baseline EPSC often exceeded the stimulation interval (10 ms) and, therefore, I_EPSC1_ under baseline and baclofen conditions was obtained from separately recorded 10–Hz trains in the same cell. In EGTA-AM experiments, 10 – 15 ml of ACSF containing 100 μM EGTA-AM were recycled continuously over the course of the application and only one cell per slice was recorded. To record optogenetically-evoked EPSCs, 200–μm thick coronal sections of ChAT-ChR2-EYFP mice were cut and light was applied via a pE-300 LED light source (CoolLED, Andover, UK) through the objective at 10 – 20 mW/cm^2^ (5 ms pulse duration).

### Stereotaxic surgery, AAV injection and calcium imaging

Adult (8 – 10 week old) ChAT-IRES-Cre mice were deeply anesthetized via i.p. injection of ketamine (90 mg/kg) and xylazine (4.5 mg/kg) followed by head fixation in the stereotaxic setup. Using a nanoliter injector (World Precision Instruments, Sarasota, FL, USA), AAV9-hSynapsin1-FLEx-axon-GCaMP6s (Addgene #112010) was injected bilaterally into the MHb at a rate of 50 nl/min for 10 min to the following coordinates (in mm from Bregma): −1.45 anterior/posterior, 0.6 lateral, −2.65 dorsal/ventral; angled at 20°. Animals were left to recover for 3 – 4 weeks and then, 1 mm thick slices were prepared and recovered as described in the electrophysiology section. After recovery, slices were transferred to the recording chamber and superfused with ACSF containing 2.5 mM Ca^2+^ at RT. Stimulating electrodes were placed bilaterally on the fasciculus retroflexus. Widefield imaging of MHb axons in the rostral/central IPN was done using a 20X 0.5NA water-immersion objective (Olympus) and a monochrome CCD camera (XM10, Olympus). To visualize the axon-GCaMP6s, blue light was emitted from a pE-300 led light source (CoolLED) through a fluorescent cube with excitation filter 460-490 nm, dichroic mirror 505 nm and barrier filter 510 nm (U-MWIB2, Olympus). GCaMP fluorescence in response to 10–Hz electrical stimulation (3–s duration, 0.2–ms pulse width, 0.5 – 2.5 V stimulation intensity) was recorded before and during the application of baclofen (1 μM). Full field-of-view frames were captured at 3 Hz and illumination was turned off in between sweeps (20–s sweep intervals).

### Calcium imaging-based modeling

For modeling, a Java-based simulator called D3D^50^, running on a Windows 10 operating system was used to calculate Ca^2+^ binding with several Ca^2+^ buffer species including GCaMP6. Assuming imaging from a whole presynaptic terminal, we allocated 16 VGCCs ^1^ in a single compartment (1 × 0.5 × 0.5 μm). In response to a single action potential (AP), each channel was allowed to carry Ca^2+^ current in a Gaussian shape. The peak amplitude was set to 0.3 pA^51,52^. The half-duration of the current was adjusted to reproduce the amplitude of the GCaMP6 fluorescent change after 10–Hz trains of 30 consecutive APs under baseline condition. The concentration of Ca^2+^-bound GCaMP6 was converted to a fluorescence change ΔF/F. We assumed that the two Ca^2+^ binding sites in GCaMP6 protein bind Ca^2+^ independently. All simulation parameters are summarized in Extended Data Table 1.

### Timed high-pressure freezing after optogenetic stimulation and freeze substitution (“Flash and Freeze”)

ChAT-ChR2-EYFP mice (8 – 10 weeks old) were deeply anesthetized, transcardially perfused and the brain was excised as described above. 200–μm thick coronal sections containing the IPN were cut and the IPN was trimmed using a razor blade. Trimmed IPN sections were left to recover as described above, followed by a 10 – 15 min incubation in ACSF containing 2.5 mM Ca^2+^ and 1 μM baclofen. Thereafter, sections were moved to the same ACSF containing 15% polyvinylpyrrolidone (PVP) followed by immediate assembly into a sapphire glass sandwich, consisting of two sapphire discs separated by a 200–μm thick spacer ring and a 400–μm thick spacer ring on the outside (Wohlwend GmbH, Switzerland), into a CLEM middle plate (Leica, Wetzlar, Germany) as described in Borges-Merjane et al. 2020^21^. After trimming, slices were never touched directly and slice transfers between solutions or into the sapphire sandwich were carried out by careful pipetting using a handcrafted, flame-polished glass pipette. The sandwich was inserted into the ICE high-pressure freezer (Leica EM) and slices were optogenetically stimulated three times at 10 Hz (3–s duration, 5–ms pulse width, 10–s interval, measured light intensity 5.9 – 10 mW/mm^2^). Slices were rapidly frozen either 100 ms (depletion group) or 10 s after the third 10–Hz stimulation (recovery group). Control group slices were not incubated in baclofen and remained unstimulated.

Freeze substitution was performed as described previously^21^. In brief, frozen sapphire sandwiches were incubated acetone containing 0.1% tannic acid at –90 °C for 20 – 24 h inside an AFS machine (Leica) under constant agitation. Thereafter, samples were washed with acetone (–90 °C) followed by incubation in acetone containing 2% osmium tetroxide and 0.2% uranyl acetate (AL-Labortechnik, Zeillern, Germany). Under continuing constant agitation, samples were then slowly warmed up to 0 °C in three steps: 1) from –90 °C to –60 °C over 2 h, incubation at –60 °C for 3 h; 2) from –60 °C to –30 °C over 4 h, incubation at –30 °C for 3 h; 3) from –30 °C to 0 over 3 h. Subsequently, slices were removed from the AFS machine, washed with acetone and propylene oxide and incubated in Durcupan resin (Sigma Aldrich; mixture of components A, B, C and D in proportion of 10:10:0.3:0.3, respectively) overnight (O/N). As slight improvement of the original Flash and Freeze method in acute slices, we performed an additional flat embedding step to precisely extract IPN subnuclei for further processing (Extended Data Fig. 3A). For flat embedding, each slice was placed on a silicon-coated glass slide, covered with an ACLAR® fluoropolymer film (Science Services, Munich, Germany) and incubated at 37 °C (1 h) followed by incubation at 60 °C (2 O/N). Using a razor blade, the rostral/central subnuclei were excised from the flat embedded slice, placed into a plastic tube (TAAB Laboratories Equipment Ltd., Aldermaston, UK) and re-embedded in Durcupan resin followed by incubation at 60 °C for 2 O/N. The resulting block was trimmed using a TRIM2 (Leica EM) and serial ultrathin sections (40 nm) were cut using a UC7 ultramicrotome (Leica EM). Finally, sections were post-stained with 2% uranyl acetate for 10 min and lead citrate for 2 min.

### “Flash and Freeze-fracture” and replica immunolabeling

Acutely cut IPN slices from ChAT-ChR2-EYFP mice were prepared, trimmed and recovered as described for “Flash and Freeze”. After recovery, slices were incubated in ACSF containing 1 mM Ca^2+^, 1 mM TEA-Cl and 100 μM 4-AP with or without 1 μM baclofen for 5 – 10 min. Thereafter, slices were moved to the same ACSF containing 2.5 mM Ca^2+^, 100 μM dynasore (HelloBio, Bristol, UK) and 15% PVP followed by assembly of the freezing sandwich consisting of a custom-made, gold-coated copper carrier, double-sided tape (150–μm thickness) and a sapphire disc. After insertion into the ICE high-pressure freezer (Leica EM), slices were stimulated twice at 10 Hz (5–ms pulse width, 3–s duration, 10–s interval), followed by a single light pulse (8–ms duration) and high-pressure freezing, timed so that the sample reached 0 °C exactly 8–ms after light stimulation onset. Light application continued for another 15 ms during the freezing process.

For freeze-fracture replication, a single metal-sapphire sandwich was inserted into an ACE900 freeze-fracture machine (Leica EM) and warmed up to –120 °C for 20 min under high vacuum of < 8 x 10^-7^ bar. Thereafter, the frozen tissue was fractured by moving the knife through the double-sided tape between the metal carrier and the sapphire disc. After fracture, carbon/platinum replication was performed as described previously^1^. Briefly, a 5-nm layer of carbon angled at 90° was evaporated onto the fracture slice, followed a 2-nm layer of platinum angled at 60° and a final 20-nm layer of carbon at 90°. After removal of the replica from the machine, slices were transferred to 0.1 M phosphate buffer (PB) containing 2% PFA for post-fixation for 1 h, followed by tissue digestion in a solution containing 2.5 % SDS, 20% sucrose in 15 mM Tris buffer (pH 8.3) at 80 °C for 18 h under gentle agitation (50 rpm).

Following the SDS treatment, immunolabeling of replicas was performed as described previously^1^. In brief, replicas were washed in washing buffer containing 0.1% Tween-20, 0.05% BSA, 0.05% NaN3 in tris-buffered saline (TBS), pH 7.4. Thereafter, replicas were incubated in the same solution containing 5% BSA (blocking buffer) for 1 h and then incubated in the same solution containing 1% BSA and primary antibodies: guinea pig anti-CAPS2 (8 μg/ml)^33^ and rabbit anti-synaptoporin (6 μg/ml; SySy, Göttingen, Germany) for 24 h. Replicas were then washed, blocked and finally incubated in secondary antibody solution (washing buffer + 5% BSA) containing 5–nm gold-conjugated anti-rabbit and 10–nm gold-conjugated anti-guinea pig antibodies (both diluted 1:30, BBI Solutions, Cardiff, UK) for 1 O/N. The next day, replicas were washed, mounted onto EM grids and dried, followed by observation in a Tecnai12 transmission electron microscope (FEI Company, Hilsboro, OR, USA) at an accelerating voltage of 120 kV.

### Immunohistochemistry

50-μm thick sections (perfused with 4% PFA) were washed in 25 mM phosphate-buffered saline (PBS) and incubated in blocking buffer containing 10% normal goat serum, 2% BSA and 0.5% triton-X 100 in 25 mM PBS for one hour, followed by O/N incubation in primary antibodies in the same buffer: anti-CAPS2 (0.5 μg/ml) or anti-synaptoporin (1 μg/ml). Slices were then washed and incubated in blocking buffer containing secondary antibodies [1:500, Alexa-488 anti-guinea pig (Molecular Probes, Eugene, OR) or Alexa-488 anti-rabbit (Molecular probes)]. Washed sections were mounted on glass slides in Mowiol (Sigma Aldrich) and observed with an LSM 800 confocal microscope (Zeiss, Oberkochen, Germany).

### Pre-embedding immunolabeling for EM

Pre-embedding immunolabeling was performed as described previously^1^. Briefly, 50–μm thick IPN slices (from 4% PFA and 0.05% glutaraldehyde perfused brains) were cryoprotected in a series of 5%, 15% and 20% sucrose in 0.1 M PB. Sections were then rapidly frozen in liquid nitrogen and immediately thawed three times. Freeze-thawed sections were treated with 50 mM glycine (Sigma Aldrich) in 50 mM TBS and were washed in TBS. Then, the sections were blocked in 2% BSA with 10% NGS in TBS followed by incubation in primary antibodies (guinea pig anti-CAPS2 (0.5 μg/ml) or rabbit anti-synaptoporin (1 μg/ml) or rabbit anti-GFP (Abcam, USA) in 2% BSA in TBS at 4 °C for 2 O/N and respective 1.4 nm gold-conjugated secondary antibodies (Nanoprobes Inc., USA) for silver intensification or in biotinylated anti-rabbit secondary antibody (Vector Labs, USA) for horseradish peroxidase (HRP) reaction for 1 ON at 4 °C. After washing in TBS and PBS, sections were post-fixed in 1% glutaraldehyde in PBS, washed with 50 mM glycine in PBS followed by another wash in PBS.

#### Silver intensification

Sections were washed in MQ water and silver intensification was performed using a commercial kit (Nanoprobes Inc., USA). Equal drops of component A (initiator) and component B (moderator) were mixed and vortexed, followed by the addition of component C (activator). After vortexing all three components properly, sections were incubated in this solution in the dark. The silver intensification reaction was stopped by adding MQ water and sections were washed with 0.1 M PB.

#### HRP reaction

Sections were washed in TBS followed by incubation in avidin-biotin complex (ABC) solution (Vector Laboratories, USA) for two hours. ABC solution was prepared 30 min prior by adding 1% avidin (A) to TBS, followed by addition of 1% biotin (B). After washing with TBS and tris-buffer (TB), sections were incubated in 0.05% di-amino benzidine (DAB) (VWR, Radnor, PA, USA) in TB for 15 min in the dark. Then, H_2_O_2_ was added to a final concentration of 0.003%. After a dark coloring became visible in the IPN, the hydrogen peroxidase reaction was stopped by adding TB. Finally, sections were washed with 0.1 M PB.

After washing sections in 0.1 M PB after silver intensification or HRP reaction, sections were fixed with 1% osmium tetroxide in 0.1 M PB for 20 min in the dark. Sections were then washed in MQ and counterstained in 1% uranyl acetate in MQ for 30 minutes in the dark. Thereafter, dehydration was done in a serial dilution of ethanol from 50%, 70%, 90%, 95% to 100%. Sections were further dehydrated in propylene oxide (Sigma Aldrich) and incubated dipped in durcupan resin O/N. On the next day, sections were flat embedded onto silicon coated glass slides and covered with an ACLAR® fluoropolymer film before polymerizing the resin for 2 O/N at 60 °C. Afterwards, rostral/central nuclei of IPN were cut by surgery blade and were re-embedded in resin in a TAAB tube. The resin was polymerized for 2 O/N at 60 °C. 70 nm sections were cut by Leica EM UC7 ultramicrotome (Leica, Germany) and were observed in a Tecnai 12 transmission electron microscope.

### Flash and Fix

200-μm thick coronal IPN slices of ChAT-ChR2-EYFP mice were prepared and recovered as described for “Flash and Freeze” experiments. After recovery, slices were incubated for 10 – 15 min in ACSF containing either 2.5 mM Ca^2+^ (“Control” and “10 Hz” groups) or 2.5 mM Ca^2+^ and 1 μM baclofen (“10 Hz + Baclofen” group). Each slice was then optogenetically stimulated three times at 10 Hz for 3 s (10 s interval) followed by immediate immersion fixation in 0.1 M PB containing 4% PFA, 0.05% glutaraldehyde and 15% picric acid under constant agitation for 45 min, followed by washing in 0.1 M PB. Control slices were immersion-fixed without light stimulation. Thereafter, pre-embedding immunolabeling and EM sample preparation was performed as described above.

### Analysis

#### Electrophysiology

All electrophysiological recordings were analyzed in Clampfit (Molecular Devices), Excel (Microsoft, Redmond, WA, USA) and Prism 8 (Graphpad, San Diego, CA, USA).

#### Flash and Freeze

Serial EM images were analyzed manually in Reconstruct software (John C. Fiala, Ph.D.). The average number of serial images analyzed per synapse was 6.3 ± 0.3 (n=52 synapses). SV diameter was measured in SVs with a clearly visible lipid bilayer as the distance between the two outer membranes through the SV center. The AZ was determined by 1) the presence of a postsynaptic density on the opposing postsynaptic side and 2) electron density inside the synaptic cleft. To minimize variability caused by different synapse sizes, distance of SVs was measured in a specified perimeter around the AZ. Specifically, a box was drawn including the AZ with the following specifications: 150 nm to the left and right of the AZ along the membrane and from there, 200 nm at right angle away from the presynaptic membrane towards the presynaptic cytosol. The open ends were connected to close the box. Inside this box, the distance of the center of all SVs to the closest inner membrane leaflet of the AZ was measured. SVs were considered docked when they had direct contact with the AZ and/or the center of the SV was within a distance of 20–nm to the inner leaflet of the presynaptic membrane.

#### Calcium imaging

Recorded image series were analyzed using Fiji for Image J. In each recording, 3 – 5 puncta were chosen as regions of interest (ROI) and fluorescence was measured at each ROI. Measured values were then normalized to the fluorescence intensity prior to the start of electrical stimulation at each ROI (F0). For each slice, the time courses of fluorescence intensity at each ROI were then averaged for baseline and baclofen groups, resulting in a single trace for each condition.

#### Flash and Fix

Ultrathin sections of chemically fixed acute slices were analyzed in Reconstruct software to count the number of silver-intensified gold particles in the AZ.

#### Flash and Freeze-fracture

Replica immunogold labeling were analyzed using Darea software^53^. In brief, presynaptic membrane profiles were manually demarcated and gold particles were automatically detected. Validity of automated detection was manually confirmed. Analysis of nearest neighbor distances (NNDs) from 10 nm particles for CAPS2 to the nearest 5 nm particle for SPO were computed to evaluate co-localization of CAPS2 and SPO. We performed Monte-Carlo fitted simulations^53^ to place randomly the SPO particles with constraints that they keep the minimum distance of 10 nm to other particles and that the distribution of NND for SPO between the simulated and original particles should not significantly differ (as assessed by two-sample Kolmogorov-Smirnov (KS) test, p > 0.1)^53^. Statistical differences in NND between CAPS2 to the original and simulated SPO particles were compared by KS test using R software (version 4.1.0).

Unless otherwise noted, all data are presented as mean ± SEM. Parametric tests were performed unless data failed Shapiro-Wilk normality test, in which case nonparametric tests were used. Statistical analysis was performed in Prism 8 unless otherwise stated. Figures were prepared using Prism 8 and Photoshop (Adobe, San Jose, CA, USA).

## Acknowledgments

We thank Erwin Neher and Ipe Ninan for critical comments on the manuscript. This project has received funding from the European Research Council (ERC) and European Commission (EC), under the European Union’s Horizon 2020 research and innovation programme (ERC grant agreement no. 694539 to Ryuichi Shigemoto and the Marie Skłodowska-Curie grant agreement no. 665385 to Cihan Önal). This study was supported by the Cooperative Study Program (e–201) of Center for Animal Resources and Collaborative Study of NINS. We thank Kohgaku Eguchi for statistical analysis, Todor Asenov from the ISTA machine shop for custom part preparations for high-pressure freezing and the electron microscopy facility at ISTA for technical support.

## Author contributions

PK performed electrophysiology experiments, calcium imaging, and high-pressure freezing of “Flash and Freeze” and “Flash and Freeze-fracture” experiments; PK and PB fractured “Flash and Freeze-fracture” samples; PB performed immunolabelings for fluorescence imaging, pre-embedding and “Flash and Freeze-fracture” replica sample preparations, and freeze substitution experiments. PB and ELM performed confocal imaging; CÖ performed viral injections, established calcium imaging and cut acute slices for calcium imaging; PK, PB and ELM performed EM imaging of “Flash and Freeze” samples; PK performed “Flash and Freeze-fracture” and “Flash and Fix” EM imaging; YN performed modeling based on calcium imaging data; CBM taught PK how to use Leica EM ICE; MS and MH made SPO KO mice using the CRISPR/Cas9 method; PJ, NB and TS provided tools and reagents. PK analyzed data and prepared figures; RS and PK conceived the study, designed experiments, and wrote the manuscript. All authors have read and jointly revised the manuscript and approved its content.

