## Supplemental Data for "A two-pool mechanism of vesicle release in medial habenula terminals underlies GABA_B_ receptor-mediated potentiation"

Author affiliations: [1]Institute of Science and Technology Austria (ISTA), 3400 Klosterneuburg, Austria; [2]Present address: Biozentrum of the University of Basel, 4056 Basel, Switzerland [3]Department of Pharmacology, Jikei University School of Medicine, Nishishinbashi, Minato-ku, Tokyo, 1058461, Japan; [4]Advanced Scientific Research Leaders Development Unit, Gunma University Graduate School of Medicine, Maebashi, Gunma 371-8511, Japan; [5]Section of Mammalian Transgenesis, National Institute for Physiological Sciences, Okazaki, 444-8585 Japan. [6]Department of Molecular Neurobiology, Max Planck Institute of Experimental Medicine, 37075 Göttingen, Germany

*These authors contributed equally

### Extended Data Figures

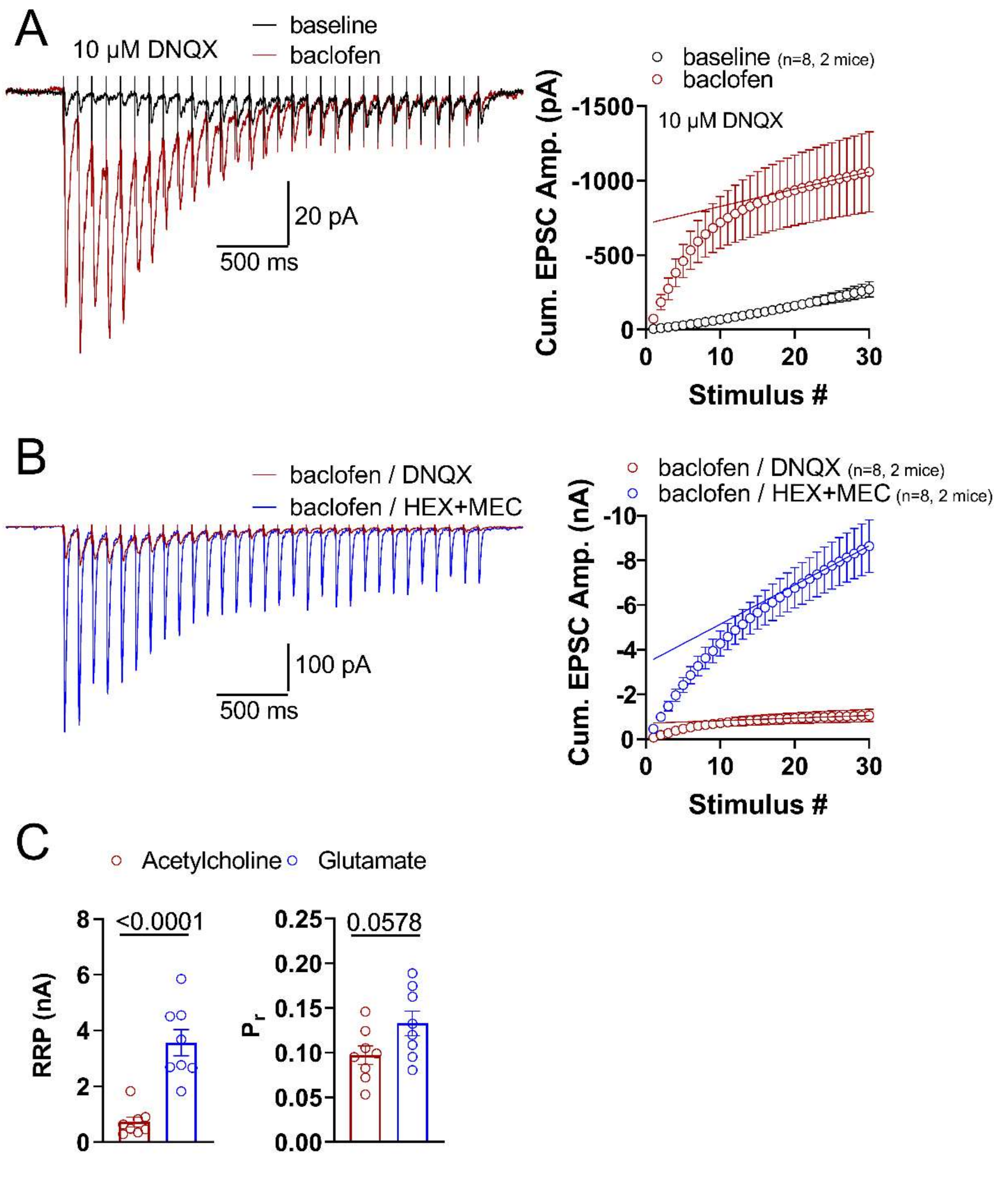

#### Extended Data Figure 1 – related to Figure 1: Comparison of cholinergic and glutamatergic phasic release

**A** Left, example traces of a 10 Hz EPSC response under baseline and baclofen conditions in the presence of AMPA receptor blocker DNQX. Right, cumulative EPSC amplitude plots of basal and baclofen EPSC trains. Linear regression line fit through the last six points (stimuli #25 – 30) of the baclofen group. **B** Left, overlay of the same cholinergic phasic release trace as in **A** with a glutamatergic phasic release trace in the presence of nicotinic acetylcholine blockers hexamethonium (HEX, 50 μM) and mecamylamine (MEC, 5 μM) from a different recording. Right, cumulative EPSC amplitude plots of cholinergic and glutamatergic phasic

EPSC trains. **C** Comparison of RRP size (left) and $P_r$ of cholinergic and glutamatergic phasic release. P values calculated by two-tailed unpaired t-test.

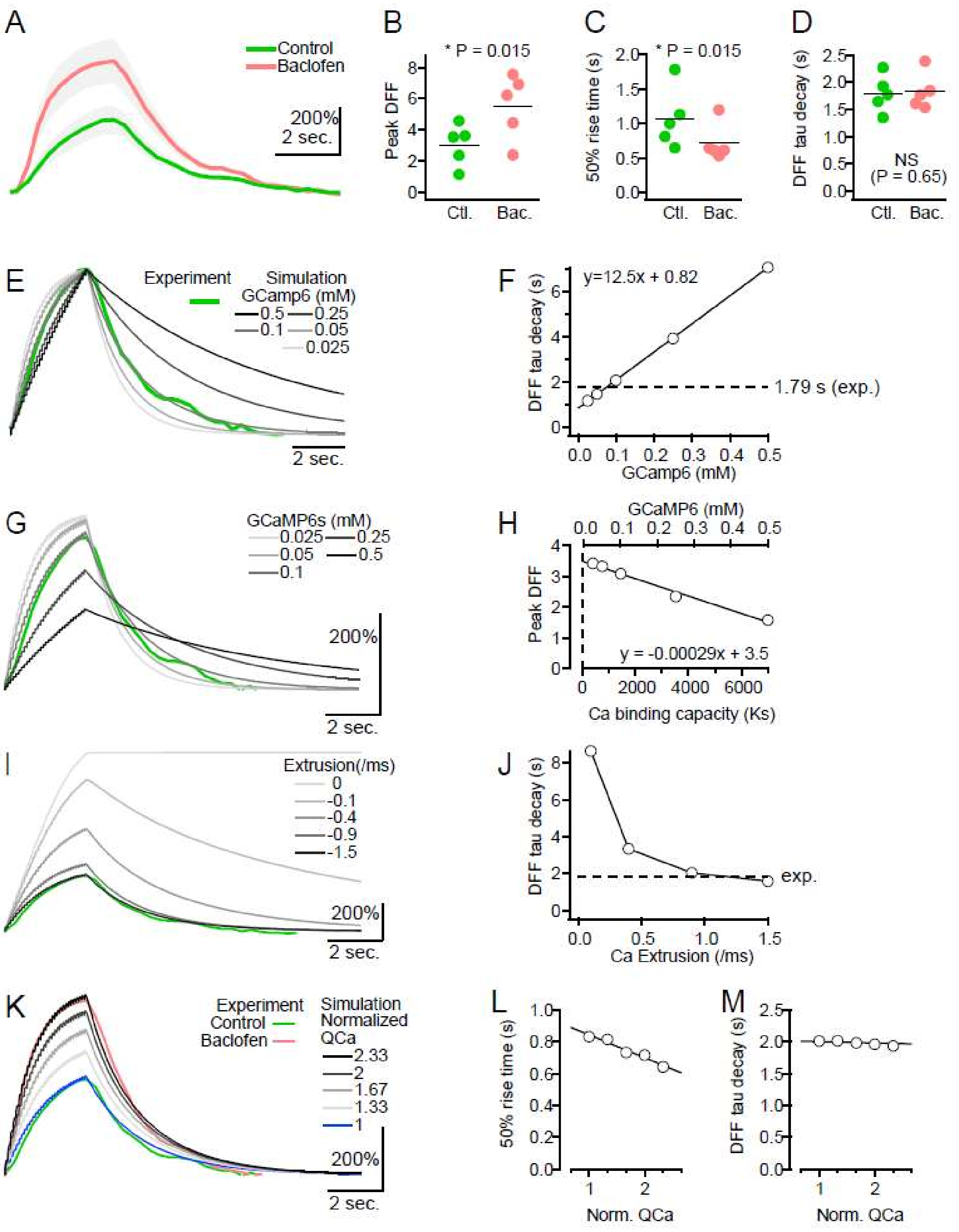

**Extended Data Figure 2 – related to Figure 2: Numerical simulations of $Ca^{2+}$ imaging in MHb terminals**

**A – D** GCaMP6 fluorescent transients induced by a 10 Hz train of 30 consecutive electrical stimuli under control condition (green) and in the presence of baclofen (pink). Each trace represents a mean and SEM (gray zone) from five recordings in the slice experiments. Peak amplitude (**B**), 50% rise time (**C**) and decay time constant (**D**) of the GCaMP6 fluorescent change in the slice experiment. **E – F** Estimation of GCaMP6 concentration in the terminal.

Because the decay time course of $Ca^{2+}$ transient in the closed single compartment model is mostly determined by the affinity and the concentration of $Ca^{2+}$ buffers, we systematically altered the GCaMP6 concentration in the simulation to examine which concentration could best reproduce the experimentally observed decay of GCaMP6. Decay time constant was linearly increased proportional to the GCaMP6 concentration. The fitting line (**F**) suggests the presence of 78 μM GCaMP6 in the MHb terminal under control condition. But for simplicity and convenience, we used 100 μM GCaMP6 in the simulations below. **G – H** The effect of $Ca^{2+}$ buffer change on the peak of $Ca^{2+}$ transients. The property of Ca buffers can affect not only the decay time course but also peak amplitude of $Ca^{2+}$ transient. Among the Ca buffer species in this simulation GCaMP6 is predominant, because other buffers (ATP and endogenous fixed buffer) have much larger Kd values thereby making a marginal contribution to total $Ca^{2+}$ binding capacity (Ks). Thus, if baclofen changes the property (affinity and concentration) of $Ca^{2+}$ buffers by some cellular mechanisms, our simulation can mimic the effect by changing GCaMP6 concentration. When we increased GCaMP6 concentration, peak ΔF/F (DFF) became smaller, the opposite direction from the observed effect of baclofen. When we decreased or even removed GCaMP6 in the simulation, the simulated DFF amplitude was increased. However, the regression line (**H**) predicted the largest DFF of 3.5, far below the observed potentiation by baclofen (**B**). **I – J** The effect of $Ca^{2+}$ extrusion on the peak of $Ca^{2+}$ transients. The other parameter affecting peak amplitude and decay time course of $Ca^{2+}$ transient in the single compartment is the $Ca^{2+}$ extrusion. In the simulation under control condition, we used extrusion rate of 0.9/ms $^{1}$. Decreasing extrusion rate achieved larger DFF like the observed effect of baclofen. This simulation also suggested that GCaMP6 was not saturated. However, reduced extrusion also prolonged the decay time course (**J**), not consistent with the imaging experiment (**D**). **K – M** The effect of $Ca^{2+}$ influx on the peak of Ca transients. In the last, we tested the possible increase in $Ca^{2+}$ influx into the MHb terminal. This can be achieved either by the change in the conductance of single $Ca^{2+}$ channel, the number of VGCCs, or the duration of $Ca^{2+}$ influx (and the combination of these). When we increased the $Ca^{2+}$ influx to 2.3-fold, the simulated traces well reproduced the GCaMP6 fluorescent change in the presence of baclofen. Increase in the $Ca^{2+}$ influx in simulation also accelerated the 50% rise time (**L**), but not significantly affected the decay time course of transients (**M**).

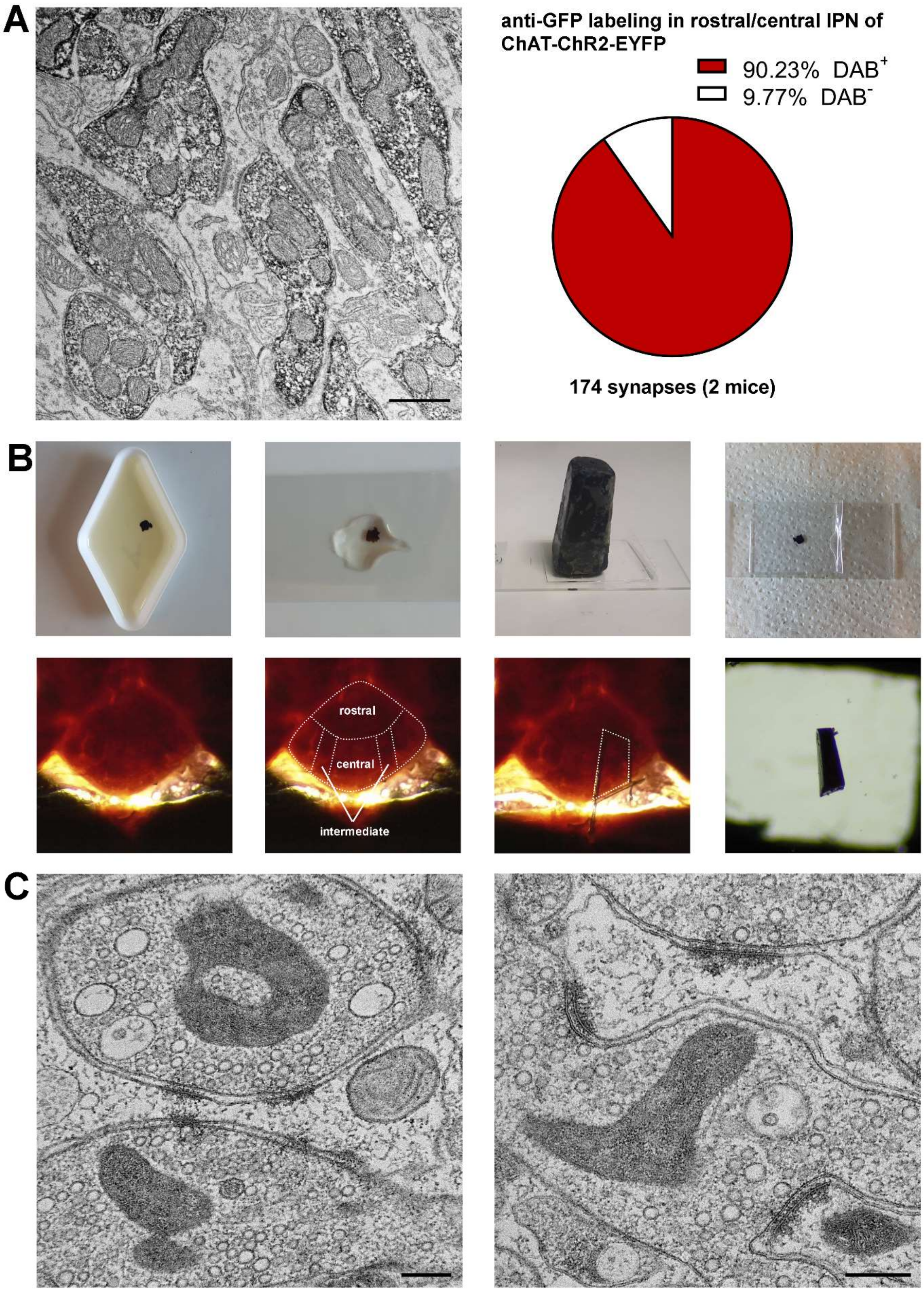

**Extended Data Figure 3 – related to Figure 4: GFP expression in MHb terminals of ChAT-ChR2-EYFP mice and "Flash and Freeze" sample preparation**

**A** Left, an example image of HRP reaction of anti-GFP labeling in ChAT-ChR2-EYFP mice in the rostral/central IPN. Three crest synapses are strongly labeled. Right, quantification of DAB-positive and negative asymmetrical synapses in the rostral/central IPN subregions. The majority of terminals was GFP-positive. **B** Processing of “Flash and Freeze” slices after freeze substitution. Top, from left to right, slices are first incubated in resin, mounted onto a glass slide, covered by an aclar film under a weight, and then flat-embedded. Bottom, from left to right, light microscopy images of a flat embedded IPN slice with demarcations of IPN subregions. A small piece containing rostra/central (including intermediate) subnuclei is trimmed. **C** Example EM images of “Flash and Freeze” samples. Left, examples of a crest synapse. Right, examples of several other asymmetrical synapses. Scale bars, 200 nm.

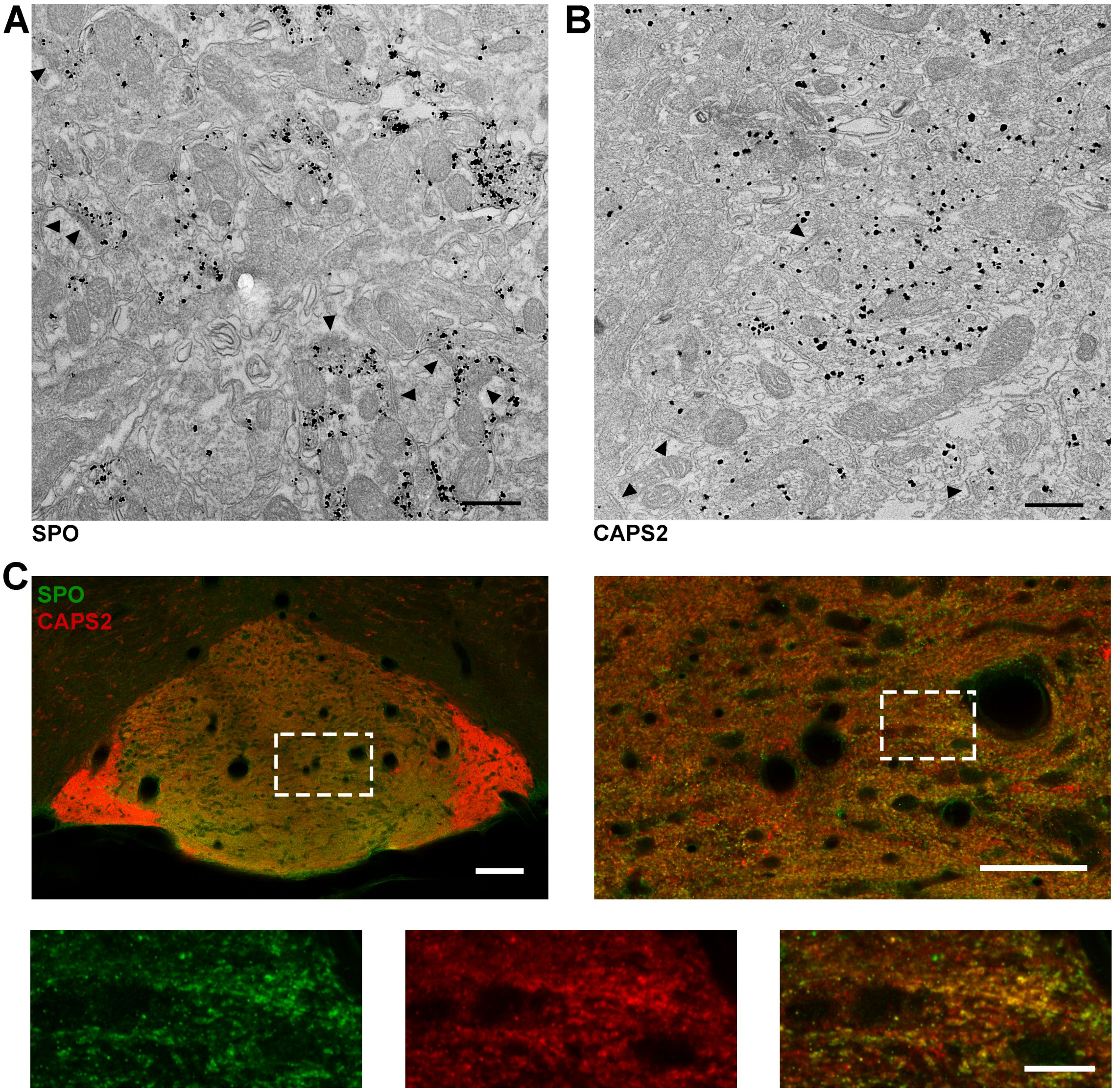

**Extended Data Figure 4 – related to Figure 5: Pre-embedding EM immunolabeling for SPO and CAPS2 in MHb terminals and confocal fluorescence imaging of SPO and CAPS2 co-labeling**

**A – B** Low magnification example images of SPO (**A**) and CAPS2 (**B**) labeling in MHb terminals. Arrow heads indicate postsynaptic density (PSD). Terminals making synapses with all of these PDSs are labeled for SPO or CAPS2. Scale bars, 500 nm. **C** Confocal fluorescent images of SPO (green) and CAPS2 (red) double immunolabeling in the whole IPN (top left, scale bar 100 μm). An area (dashed box) on the left is shown on the right at higher magnification (top right, scale bar 50 μm). An area (dashed box) on the right is further magnified (bottom) to show individual SPO- (green) and CAPS2-positive (red) puncta. Overlay shown on the right. Scale bar, 10 μm.

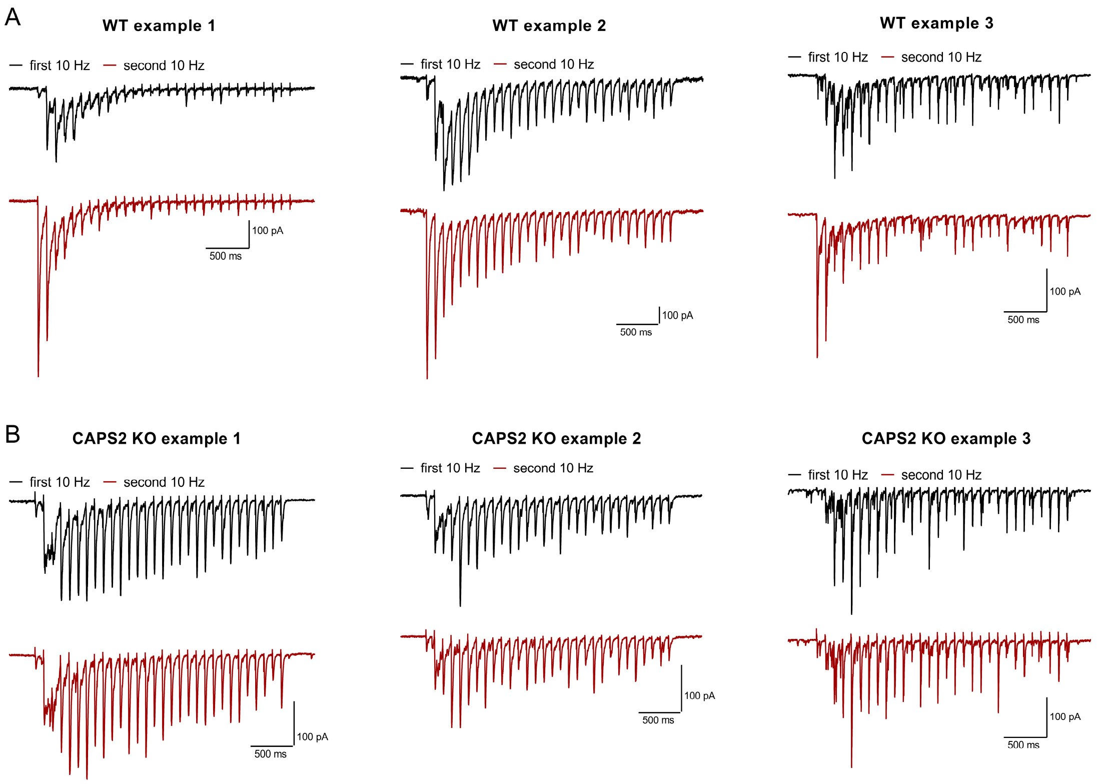

**Extended Data Figure 5 – related to Figure 5: Comparison of phasic EPSC responses to the first and second 10 Hz stimulation in WT and CAPS2 KO mice**

Example traces from three recordings of 10 Hz stimulation trains in the presence of baclofen at the first and second stimulation (20–s interval) in WT (**A**) and CAPS2 KO mice (**B**).

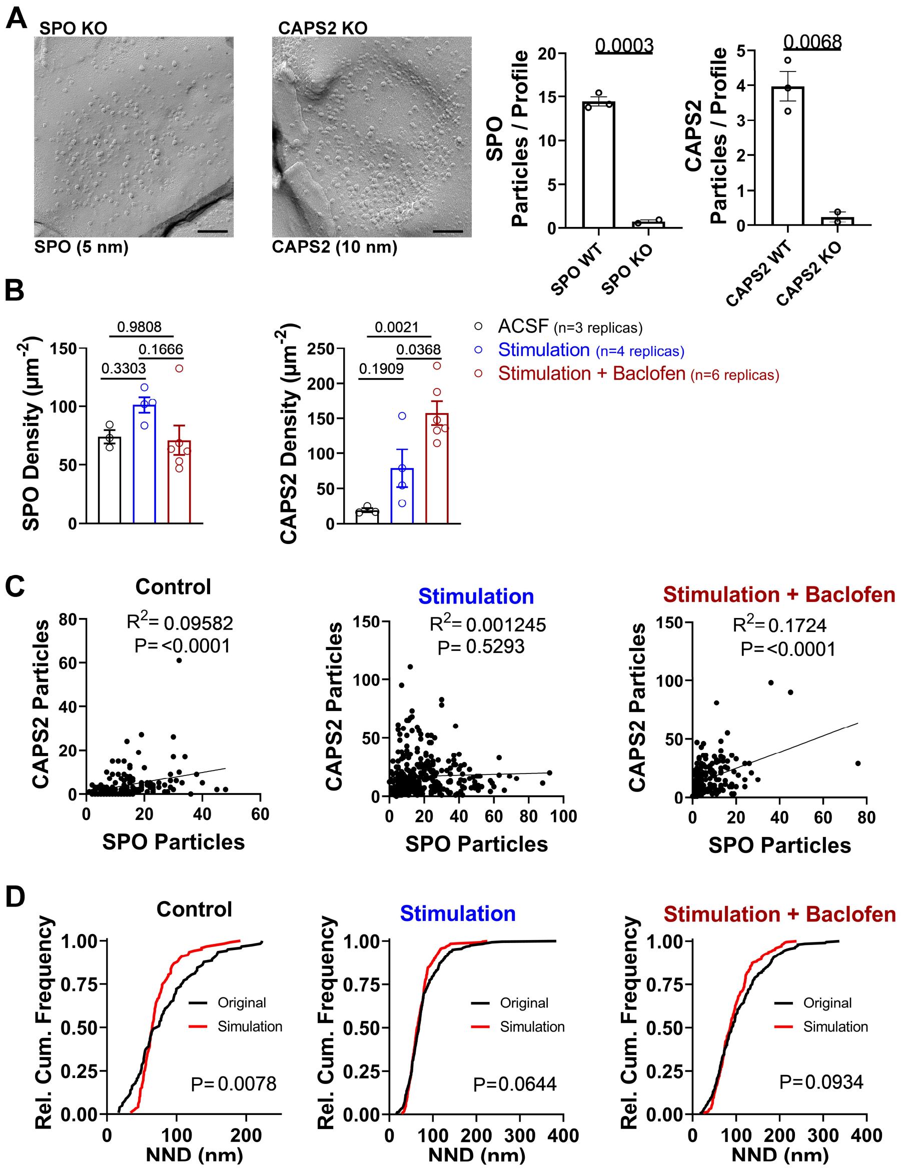

**Extended Data Figure 6 – related to Figure 8: Quantification of antibody specificity and additional "Flash and Freeze-fracture replica" analyses**

**A** Left, example images of SPO and CAPS2 labeling in acute slice replicas of the respective KO mice. Right, quantification of SPO and CAPS2 particle numbers per profile in replicas of WT and corresponding KO mice. P value calculated by two-tailed unpaired t-test. Scale bars, 100 nm **B** Quantification of SPO (left) and CAPS2 (right) particle densities in presynaptic active zones in three conditions, control (ACSF, without stimulation and baclofen), stimulation in the absence and presence of baclofen. P values calculated by one-way ANOVA with Tukey post hoc test. **C** Correlation analysis of SPO and CAPS2 particle numbers. $R^2$ and P values calculated by Pearson correlation. **D** Comparison or nearest neighbor distances (NNDs) from CAPS2 (10–nm gold particles) to SPO (5–nm gold particles) between real data (Original) and simulated random particle distributions (Simulation). P values derived from Kolmogorov-Smirnov test.

### Extended Data Tables

| Simulation Parameters | value | units | Reference |
|---|---|---|---|
| Simulation voxel size | 0.1 | µm | |
| Simulation volume (x,y and z) | 1 × 0.5 × 0.5 | µm | |
| Time step for simulation | 0.38 | µs | |
| Output | 1k | Hz | |
| ***$Ca^{2+}$ entry per single AP*** | | | |
| Maximal single channel current | 0.3 | pA | Calculated from Sheng et al. 2012 [2] |
| Half-duration (Gaussian shape) | 0.19 | ms | Nakamura et al., 2015 [3] |
| Size of $Ca^{2+}$ entry site | 0.2 × 0.2 | µm | Bhandari et a., 2021 [4] |
| Number of open channels | 6 | | Bhandari et a., 2021 [4] |
| ***$Ca^{2+}$*** | | | |
| Diffusion coefficient | 0.22 | $\mu m^2\ ms^{-1}$ | Allbritton et al, 1992 [5] |
| Resting free $Ca^{2+}$ concentration | 10 | nM | |
| $Ca^{2+}$ extrusion | -0.9 | $ms^{-1}$ | Helmchen et al, 1997 [1] |
| *Endogenous fixed buffer properties* | | | |
| $k_{on}$ | 100 | $mM^{-1}ms^{-1}$ | Nakamura et al., 2015 [3] |
| $k_{off}$ | 10 | $ms^{-1}$ | |
| Concentration | 4.0 | mM | Calculated from Helmchen et al, 1997 [1] |
| ***ATP $Ca^{2+}$ binding properties*** | | | |
| $k_{on}$ | 500 | $mM^{-1}ms^{-1}$ | Naraghi & Neher, 1997 [6] |
| $k_{off}$ | 100 | $ms^{-1}$ | |
| Diffusion coefficient | 0.22 | $\mu m^2\ ms^{-1}$ | |
| Concentration | 0.2 | mM | |
| ***GCaMP6s properties*** | | | |
| $k_{off}$ | 0.00112 | $ms^{-1}$ | Chen et al., 2013 [7] |
| Kd | 144 | nM | |
| $R_{max}/R_{min}$ | 63.2 | | |
| Concentration | 0.1 | mM | Adjust to experiments |
| Diffusion coefficient | 0.02 | $\mu m^2\ ms^{-1}$ | Adopted from CR, Faas et al, 2007 [8] |

**Extended Data Table 1 – related to Extended Data Figure 2: Parameters for simulations of fluorescent $Ca^{2+}$ transients**
